# Rab11 regulates autophagy at dendritic spines in an mTOR- and NMDA-dependent manner

**DOI:** 10.1101/2022.05.20.492668

**Authors:** Aleksandra Janusz-Kamińska, Agnieszka Brzozowska, Aleksandra Tempes, Malgorzata Urbanska, Magdalena Blazejczyk, Jacek Miłek, Juan Zeng, Katarzyna Kisielewska, Jacek Jaworski

**Affiliations:** Laboratory of Molecular and Cellular Neurobiology, International Institute of Molecular and Cell Biology, Warsaw, Poland; Laboratory of Molecular Basis of Synaptic Plasticity, Centre of New Technologies, University of Warsaw

**Keywords:** Atg9A, Rab11, mTOR, autophagy, autophagosomes, neurons, synaptic plasticity

## Abstract

Synaptic plasticity is a process that shapes neuronal connections during neurodevelopment, learning, and memory. Autophagy is a mechanism that allows cells to degrade their unnecessary or dysfunctional components. Autophagosomes appear at dendritic spines in response to plasticity-inducing stimuli. Autophagy defects contribute to altered dendritic spine development, autistic-like behavior in mice, and neurological disease. While several studies explored the involvement of autophagy in synaptic plasticity, the steps preceding autophagosome emergence at the postsynapse remain unknown. Here we show a postsynaptic association of autophagy-related protein 9A (Atg9A), known to be involved in the initial stages of autophagosome formation, with Rab11, a small GTPase that regulates endosomal trafficking. Rab11 activity is necessary for the maintenance of Atg9A-positive structures at dendritic spines. Inhibition of mTOR increased Rab11 and Atg9A interaction and increased the emergence of autophagosomes in dendritic spines when coupled to NMDA receptor stimulation. Dendritic spines with newly formed autophagosomes were more resistant to NMDA-induced morphologic change. These results collectively suggest that autophagy initiation in dendritic spines depends on an activity-dependent Rab11a-Atg9A interaction regulated by mTOR.

## Introduction

Macroautophagy or autophagy is sequestering and degrading proteins or even whole organelles in bulk (Yin et al., 2016). Mammalian/mechanistic target of rapamycin (mTOR) kinase regulates autophagy in mammalian cells (Noda and Ohsumi, 1998, Jung et al., 2009, Ganley et al., 2009). Autophagosomes are double-membrane structures formed during autophagy that fuse with lysosomes to digest autophagolysosome contents (Yin et al., 2016).

Autophagy plays an essential role in neuronal development and synaptic function, which can be altered in neurological disease (Stavoe and Holzbaur, 2019). Long-term mTOR inhibition increases the number of autophagosomes in the neuronal soma of the mouse cortex (Mizushima et al., 2004; Boland et al., 2008). Autophagy occurs continuously in axons (Maday et al., 2012, Maday and Holzbaur, 2016) and contributes to synaptic plasticity – defined as a process that changes the strength of synaptic connections (Bosch et al., 2012, Lai and Ip, 2013, Borczyk et al., 2019). Silencing or enhancing global synaptic activity in cultured neurons alters the movements of autophagosomes in dendritic shafts (Kulkarni et al., 2021). One form of synaptic plasticity is long-term depression (LTD), defined as the prolonged weakening of the strength of synaptic connections (Malenka and Bear, 2004). Currently, several studies report the connection of LTD to neuronal autophagy. Autophagosomes in neuronal soma, dendrites, and dendritic spines are at low levels under basal conditions and increase in number upon the induction of NMDA-cLTD (Shehata et al., 2012). Finally, autophagosomes appear in dendrites shortly after NMDA-cLTD treatment (Kallergi et al., 2022). Postsynaptic scaffold proteins and glutamate receptors degrade via autophagy, which modulates synaptic plasticity (Nikotelopulou et al., 2017, Compans et al., 2021, Kallergi et al., 2022). However, the origin of autophagy at the postsynapse remains unknown.

Studies in non-neuronal cells provide insight into the mechanism of autophagy initiation. Other proteins, such as Unc-51 like autophagy activating kinase 1 (Ulk1), autophagy-related protein (Atg) 5, Atg13, and other Atg family proteins, participate in autophagy initiation (Yang and Klionsky, 2010). In neurons, Ulk1 puncta increased in the dendrites upon NMDA-cLTD treatment (Kallergi et al., 2022). In axons, autophagosomes emerge from endoplasmic reticulum domains positive for FYVE-containing protein 1 and mature during retrograde axonal transport (Maday et al., 2012, Maday and Holzbaur, 2014).

Recent studies report that Rab11, a guanosine triphosphatase (GTPase), is associated with recycling endosomes in this process in non-neuronal cells and recycling endosomes act as a membrane source for autophagosomes (Longatti et al., 2012, Welz et al., 2014). Rab11-positive recycling endosomes colocalized with Ulk1 and Atg9A, dispatching the membrane, causing the emergence of autophagosomes (Welz et al., 2014). Atg9A is essential for mammalian autophagy. It interacts with the autophagy initiation site (Orsi et al., 2012) and is known to traffic through recycling endosomes (Imai et al., 2016). Puri et al. (2018) proposed that Rab11-positive recycling endosomes in HeLa cells form a platform for assembling autophagy machinery and autophagosome formation.

Much remains unknown about the steps preceding the phagophore formation via the Ulk1 complex and the emergence of autophagosomes at the dendritic spines, morphologically and functionally distinct compartments crucial for neuroplasticity, memory, and learning. Here we demonstrate a novel role for Rab11 in the initiation of autophagy at the postsynapse. Rab11-positive vesicles are present in dendritic spines, and their mobility is reduced upon treatment with the mTOR inhibitor INK128. Atg9A is highly prevalent at the dendritic spines, which was not previously known, and Rab11 is necessary for the postsynaptic accumulation of Atg9A. Rab11 co-immunoprecipitates with Atg9A and microtubule-associated protein 1A/1B-light chain 3 (LC3), an autophagosome membrane associated protein marker. Live observation of the same neurons over time revealed the dynamics of postsynaptic autophagosome formation upon mTOR inhibition and NMDA stimulation, which corresponds in a spatiotemporal manner to Rab11 and Atg9A accumulation and association.

## Significance statement

Autophagy, loosely translated as self-eating, is a crucial process in neurons, recently shown to be involved in neuronal connectivity and synaptic strength changes known as LTD (long-term depression) and LTP (long-term potentiation). These changes, or neuroplasticity, underlie learning, memory, and neurodevelopment. However, the early steps of autophagy in postsynaptic compartment neurons are not clear. Here, we show that the interaction between Rab11a GTPase and Atg9A proteins depend on the mTOR kinase and precedes autophagy at the synapses, thus likely participating in neuroplasticity.

## Methods

### Primary hippocampal cell cultures, treatments, and transfections

The animals used to obtain neurons for further experiments were sacrificed according to protocols that followed the European Community Council Directive 2010/63/EU. Rat embryonic primary hippocampal cell cultures were prepared and transfected using Lipofectamine 2000 (Thermo Fisher Scientific, Invitrogen, catalog no. 11668019) as described previously (Janusz et al., 2013, Jaworski et al., 2005). DIV21-25 neurons with mature, stabilized dendritic spines were used for all the experiments. INK128 (300 nM; Cayman Chemical, catalog no. 11811-5) was used for mTOR inhibition. Chemical LTD was induced by 3 min of NMDA (50 µM; Tocris Biosciences, catalog no. 0114/50) stimulation and washout (Shehata et al., 2012). The neurons were then imaged for up to 60 min in live experiments or fixed after 25 min.

### Antibodies

For Western blot (WB), immunofluorescence (IF), and immunoprecipitation (IP), the following primary antibodies were used: rabbit anti-phospho-Akt (Ser473; Cell Signaling Technology, catalog no. 4060, 1:1000 for WB), mouse anti-Akt (Cell Signaling, catalog no. 2920, 1:1000 for WB), rabbit anti-Atg9A antibody (Thermo Fisher Scientific, catalog no. PA5-21043, 1:200 for IF, 1:100 for WB), rabbit anti-caspase 3 antibody (Cell Signaling, catalog no. 9662; 1:1000 for WB), rabbit anti-phospho-GluA1 (Ser845; Millipore, catalog no. 04-1073, 1:500 for WB), mouse anti-GluA1 (Santa Cruz Biotechnology, catalog no. sc-55509, 1:100 for WB), rabbit anti-Hook3 antibody (Proteintech, catalog no. 15457-1-AP, 1:100 for IF), rabbit anti-LC3B antibody (Thermo Fisher Scientific, catalog no. PA1-46286, 1:200 for WB), rabbit anti-LC3B antibody (Sigma Aldrich, catalog no. L7543, 1:2000 for WB), rabbit anti-SQSTM1/p62 polyclonal antibody (Cell Signaling, catalog no. 9662; 1:1000 for WB), rabbit anti-syntaxin 12 antibody (Proteintech, catalog no. 14259-1-AP, 1:100 for IF), mouse anti-Rab11 antibody (BD Biosciences, catalog no. 610656, Clone 47/Rab11, 1:50 for IF), rabbit anti-Rab11 antibody (Cell Signaling Technology, catalog no. mAB 5589, 1:100 for IF), and rabbit anti-Rab11a antibody (Thermo Fisher Scientific, catalog no. 71-5300, 5 µg for IP), rabbit anti-phospho ribosomal protein S6 (Ser235/236; Cell Signaling Technology, catalog no. 4858, 1:1000 for WB), mouse anti-ribosomal protein S6 (Cell Signaling Technology, catalog no. 2317, 1:1000 for WB), mouse anti-α-tubulin (Sigma-Aldrich, catalog no. T5168, 1:5000 for WB). Although all our Rab11 antibodies were raised against Rab11a synthetic epitopes, they could also react with Rab11b. In the present study, we did not attempt to resolve this distinction. The following secondary antibodies were used for immunofluorescence and Western blot: anti-rabbit Alexa Fluor 405 (Thermo Fisher Scientific, catalog no. A-31556), anti-rabbit Alexa Fluor 488 (Thermo Fisher Scientific, catalog no. A-11034), anti-rabbit Alexa Fluor 647 (Thermo Fisher Scientific, catalog no. A-31573), and anti-mouse Alexa Fluor 555 (Thermo Fisher Scientific, catalog no. A-31570). Peroxidase AffiniPure donkey anti-rabbit IgG (H+L) (Jackson ImmunoResearch; catalog no. 711-035-152), peroxidase AffiniPure donkey anti-mouse IgG (H+L) (Jackson ImmunoResearch; catalog no. 715-035-150), IRDye 680RD donkey anti-Mouse IgG (LiCOR Biosystems cat. no. 926-68072), IRDye 680RD donkey anti-Rabbit IgG (LiCOR Biosystems cat. no. 926-68073), IRDye 800CW donkey anti-Mouse IgG (LiCOR Biosystems cat. no. 926-32212), IRDye 800CW donkey anti-Rabbit IgG (LiCOR Biosystems cat. no. 926-32213) and VeriBlot for immunoprecipitation detection reagent (HRP) (Abcam; catalog no. ab131366). Secondary antibodies for Western blot were diluted at either 1:5000 (HRP-conjugated) or 1:10000 (IRDye-conjugated).

### Plasmids

The following plasmids were used for the transfection of cultured cells: pMXs-IP-EGFP-LC3 (Hara et al., 2008) (gift from Noboru Mizushima; Addgene plasmid no. 38195; http://n2t.net/addgene:38195; RRID: Addgene_38195), pMXs-puro-RFP-ATG9A (Koyama-Honda et al., 2013), (gift from Noboru Mizushima; Addgene plasmid no. 60609; http://n2t.net/addgene:60609; RRID: Addgene_60609), piRFP702-N1 (Secherbakova and Verkhusha, 2013) (gift from Vladislav Verkhusha; Addgene plasmid no. 45456; http://n2t.net/addgene:45456; RRID: Addgene_45456), pmScarlet-i_C1 (Bindels et al., 2017) (gift from Dorus Gadella; Addgene plasmid no. 85044; http://n2t.net/addgene:85044; RRID: Addgene_85044), mRFP-Rab11a, and GW1-GFP-Rab11a (Esteves da Silva et al., 2015) (gift from Casper Hoogenraad). GW1-GFP-Rab11S25N and Rab11S20V were obtained by QuickChange mutagenesis using the following primers: Rab11S25N (5’- GAGATTCTGGTGTTGGAAAGAATAATCTCCTGTCTCGATTTAC-3’ and 5’- GTAAATCGAGACAGGAGATTATTCTTTCCAACACCAGAATCTC-3’), Rab11S20V (5’-ACTACCTCTTTAAAGTTGTCCTTATTGGAGATGTTGGTGTTGGAAAGAGTAAT-3’ and 5’- ATTACTCTTTCCAACACCAACATCTCCAATAAGGACAACTTTAAAGAGGTAGT-3’). Plasmid pmScarlet-i-Atg9A was prepared by the sequence- and ligation-independent cloning (SLIC) (Li et al., 2012) of Atg9A cDNA from pMXs-puro-RFP-ATG9A to pmScarlet-i_C1. The plasmid containing the insert was digested with BsrGI, and the backbone was digested with XhoI. The SLIC primers were the following: 5’- GTACAAGTCCGGACTCAGATCTCGAATGGCGCAGTTTGACACTG-3’ and 5’- TGCAGAATTCGAAGCTTGAGCTCGAACGCGTGAATTCGTTCTAGAAAC-3’.

### Imaging of live neurons

For the Rab11 assays, neurons were cultured on MatTek multiwell glass-bottom plates (P12G-1.0-14-F), treated with 1 M HCl for 15 min, rinsed three times with phosphate-buffered saline (PBS), rinsed three times with distilled H_2_O, and coated with poly-D-lysine and laminin (Jaworski et al., 2005). For the autophagy assays, neurons were cultured on 18 mm diameter glass slides in 12-well plates prepared and coated according to routine laboratory protocols (Jaworski et al., 2005).

Imaging occurred using the Andor Revolutions XD spinning disc microscope with 63x lens and 1.6 Optovar at 1004 x 1002-pixel resolution, equipped with a double incubation chamber with an external incubation cage and a heating insert with CO_2_ delivery. The CO_2_ valve was opened at least an hour before imaging, and the setup was warmed to 37°C overnight. In the case of MatTek multiwell glass-bottom plates, cells were imaged in the same culture media they were cultured in (Neurobasal/B27 Supplement/Glutamax, all from Thermo Fisher Scientific). In the case of neurons cultured on the coverslips, Chamlide magnetic chamber (Quorum Technologies Inc.) was used with 500 µl of the culture media from the same well. All subsequent media changes and washes were conducted using warm culture media from the same plate. The temperature was calibrated in the incubation cage and the sample before the experiments. Both temperature and CO_2_ flow were monitored in the incubation cage throughout the experiment. Cells were equilibrated for 30 minutes on the microscope stage and monitored for any changes before imaging.

For the Rab11 mobility assay, neurons were transfected with plasmids encoding RFP and GFP-Rab11a. The next day, time-lapse live imaging occurred using an Andor confocal spinning disc microscope at 300 ms per frame (63x objective, 1.4 NA, Optovar magnification changer 1.6, calibrated pixel size = 68 nm). Imaging began no later than 18 h after transfection. Neurons were recorded for 1 min at the start of the experiment (preincubation), after 30 min, and then for the third time after 20 min after INK128 (300 nM) or mock treatment. All dendritic spines in the field of view were marked with separate regions of interest (ROIs) in ImageJ software. Each spine was analyzed for the presence of Rab11a vesicles. Briefly, vesicles that entered or left the spine or at least reached the base of the spine neck were classified as mobile. Immobile vesicles remained in the spine head and did not travel. Dendritic spines were classified into spines that contained mobile or immobile vesicles. Spines that harbored both types were classified as spines that contained mobile vesicles. The same dendritic spines were analyzed at -30 min (preincubation), 0 min, and 20 min time points.

For the Rab11a and Atg9A assays, neurons were transfected with plasmids encoding either GFP-Rab11a or its mutants (DN GFP-Rab11 and CA GFP-Rab11), supplemented with ScarletI-Atg9A and iRFP702 to visualize dendrites and dendritic spines. The next day, three-channel snapshots were taken at the dendritic spine level. Dendritic spines that were in focus were marked as ROIs and counted. All ROIs were converted into a mask. Masked channels were then subjected to object-based colocalization in ImageJ using the JACoP plugin (Bolte and Cordelières, 2006).

For the autophagy assay, neurons were transfected with plasmids encoding EGFP-LC3 and Scarlet-I to visualize the cell body, dendrites, and spines. The next day, cells were pre-incubated with INK128 (300 nM) for 15 min or kept without treatment for 15 min. NMDA (50 µM) was added for 3 min, then the medium was aspirated and replaced with either conditioned medium or conditioned medium supplemented with INK128 (300 nM). Imaging began 3 min after medium replacement (i.e., 6 min after stimulation). For all four variants (control, INK128 only, NMDA only, and INK128+NMDA), full-stack images (Z-step = 0.6 µm) were taken at 2-min intervals before stimulation 40 min after stimulation or, in the case of the control, 40 min after first imaging. Z-stacks were processed as maximum Z projections using Andor software. For the analysis, neurons with high EGFP-LC3 expression, neurons with aberrant dendritic spine morphology, or neurons that degraded over time were rejected. Neurons were analyzed using ImageJ to determine the number of dendritic spines and new puncta that appeared after stimulation or in the equivalent period.

Neurons were subjected to the INK128 preincubation and subsequent NMDA stimulation in the presence of INK128, as described above, to analyze dendritic spine morphology. Images were collected every 2 min for 10 minutes before and after the stimulation. Imaging concluded after at least 46 min following the stimulation. The last frame taken before the stimulation (baseline) and the last frame at 46 min after the stimulation were used to analyze dendritic spine length. Spine length was measured with line ROI for each visible spine in the Scarlet-I channel. Dendritic spines before and after stimulation were compared. Those with more than 25% change in length (either by elongation or shrinking) were arbitrarily classified as those that changed shape.

Afterward, the images were analyzed in the EGFP-LC3 channel. During the movie, post-stimulation, dendritic spines that did not exhibit any autophagic puncta were classified as spines with no autophagosomes. During the movie post-stimulation, dendritic spines that exhibited autophagosome puncta were classified as spines with autophagosomes. Dendritic spines possessing autophagosome puncta at baseline were not counted. Finally, we correlated the appearance of autophagosomes after INK128+NMDA treatment with the change of dendritic spine shape and presented the result as a percentage of spine shape change in two categories, autophagosome negative or positive, in each cell.

### Immunofluorescence and super-resolution Airyscan imaging

Primary rat hippocampal neurons were cultured on glass coverslips in 24-well plates, treated as described above, and fixed for 15 min with 4% paraformaldehyde, followed by 2 min of -20°C methanol post-fixation to retrieve membrane epitopes. The glass coverslips were then rinsed in PBS and blocked for 30 min in immunofluorescence (IF) solution 1 (0.1% saponin, 0.2% gelatin, and 5 mg/ml bovine serum albumin in PBS), followed by a single wash and subsequent incubation in IF solution 2 (0.2% gelatin and 0.01% saponin in PBS) with the primary antibody in a cold room overnight. The coverslips were then rinsed three times in IF solution 2 and incubated in IF solution 2 with a secondary fluorescent antibody. The coverslips were then rinsed three times in PBS and mounted in Prolong Gold antifade reagent (Thermo Fisher Scientific, catalog no. P10144). The coverslips were imaged on a Zeiss LSM 800 microscope with the Airyscan module (63x objective, 1.4 NA, 1.3⋅ digital magnification) and then processed using the Airyscan module at default settings. The final image resolution was 28 nm/pixel. For colocalization, the GFP channel was masked to analyze only areas that contained dendritic spines of all shapes, including neck and head spines. Spines within approximately 10 µm to the cell soma were omitted. Object-based colocalization was then performed based on centers of mass (Rab11 channel) - particles coincidence using the JACoP plugin (Bolte and Cordelières, 2006) and two focal planes that contained dendritic spines that were in focus out of five focal planes that encompassed the whole neuron.

### Synaptoneurosome isolation, treatment, and protein co-immunoprecipitation

Cortical and hippocampal synaptoneurosomes from the brain of adult mice were prepared according to our previously published protocol (Janusz et al., 2013). Final extracts were diluted in 500 µl synaptoneurosome buffer (125 mM NaCl, 1.2 mM MgSO4, 2.5 mM CaCl2, 1.53 mM KH2PO4, 212.7 mM glucose, 4 mM NaHCO3, pH 7.4, set with carbogen, supplemented with Protease Inhibitor Cocktail [Sigma-Aldrich] and 100 U/ml mammalian placental RNase inhibitor [Fermentas]), per brain, pooled, and divided into 250 µl aliquots. Fresh synaptoneurosomes were then incubated at 37°C for 15 min with INK128 (600 nM) or without and then stimulated for 5 min with NMDA (50 µM). After 5 min, all the variants were diluted 5x with a preheated synaptoneurosome buffer to reduce the concentration of NMDA under the threshold known to induce cLTD (Lee et al., 1998) with or without INK128 (600 nM) and incubated for 25 min. After this time, samples were either lysed for WB analysis of GluA1 and S6 phosphorylation level or were centrifuged at 1000x g for 15 min, resuspended in immunoprecipitation buffer (10 mM HEPES, 1.5 mM MgCl2, 2 mM EDTA, 0.6% CHAPS, 120 mM NaCl, and protease inhibitors [benzamidine, aprotinin, Pefabloc; Sigma-Aldrich]), and allowed to incubate on ice for 30-60 min. Extracts were precleared with Dynabeads G (Invitrogen) and precipitated with 5 µg Rab11 antibody or total rabbit IgG. Finally, the beads were washed 3-5 times with immunoprecipitation buffer, and protein was eluted with Laemmli buffer at 98°C for 10 min. Protein samples were separated with sodium dodecyl sulfate-polyacrylamide gel electrophoresis (SDS-PAGE), followed by Western blot (WB) analysis.

### Western blot

SDS-PAGE occurred in gradient gels (12%/15%, 1 mm thick) to analyze IP results. The gels were run at 120 V for approx. 1.5h until the dye reached the bottom of the gel. Then, the gels were wet transferred at 400 mA for 90 min to methanol-activated 0.2 µm pore size PVDF membrane (Bio-Rad). Blots were blocked in 5% nonfat milk in Tris-buffered saline (0.5 mM Tris-HCl [pH 7.5] and 1.5 mM NaCl) with 0.1% Tween-20 (TBST) for 1 h and then incubated overnight in a cold room in primary antibody diluted in 5% milk in TBST. Following incubation, they were washed three times for 10 min each in TBST and then incubated for 1.5-2 h in HRP-conjugated secondary antibody diluted in 2.5% milk in TBST. Finally, blots were developed for ∼2-3 min in a freshly made chemiluminescence developing buffer (100 mM Tris [pH 8.5], 1.25 mM 3-aminoftalhydraside, 200 μM coumaric acid, and 0.01% hydrogen peroxide). Signals were detected on radiographic film after digital processing, and band intensities were measured as mean gray values using Image J software. Mean gray values of Atg9A and LC3 were normalized to mean gray values of IgG as an external loading control and then to the control (untreated) immunoprecipitation sample. Mean gray values of every other sample are presented as ratios. Quantitative analysis of S6, Akt, and GluA1 phosphorylation levels was performed using the Infrared Odyssey Imaging System (LI-COR Biosciences) described previously (Urbanska et al., 2018, Urbanska et al., 2019).

### Statistical analysis

The number of cells and culture batches analyzed and a description of statistical tests used are provided in the figure legends. All the analyses employed GraphPad Prism 8 software.

## Results

### Rab11a mobility decreases at dendritic spines upon mTOR inhibition

An earlier study showed that Rab11-positive recycling endosomes contribute to autophagy (Longatti et al., 2012). A lack of nutrients inhibits mTOR complex 1 (mTORC1), one of two protein complexes formed by mTOR, and the suppression of mTORC1 activity induces autophagy, which was demonstrated in both yeast and non-neuronal cells (Noda and Ohsumi, 1998, Levine et al., 2004). mTORC1 is needed to maintain the transferrin receptor recycling through the endosomal pathway in mouse embryonic fibroblasts (Dauner et al., 2017). Considering these findings in non-neuronal cells, we performed a Rab11 mobility assay to investigate the influence of mTOR kinase on Rab11a at dendritic spines. To suppress mTOR, we applied INK128, an ATP-competitive inhibitor of mTORC1 and mTORC2. mTORC1 is an inhibitor of autophagy, and mTORC2 likely influences autophagy through Akt activation (Mammucari et al., 2007), protein kinase Cα/β (Renna et al., 2013), or changes in mitochondrial homeostasis (Aspernig et al., 2019). In our experiments, we used embryonic rat hippocampal neurons that had mature dendritic spines (day *in vitro* [DIV] 21-25) with a gentle lipofectamine-based transfection protocol suited to older neurons which allows them to be transfected without showing signs of cytotoxicity (**Fig 1A**). For the Rab11 mobility assay, we transfected neurons with plasmids encoding green fluorescent protein (GFP)-Rab11a and red fluorescent protein (RFP) to visualize transfected neurons’ morphology (including dendrites and dendritic spines). We performed time-lapse imaging of transfected cells using a confocal spinning-disc microscope the next day. We imaged each neuron three times: at baseline (preincubation; not shown), after 30 min of incubation on the microscope stage to monitor for potential signs of neuronal toxicity and any changes in imaging conditions (e.g., temperature; 0 min. timepoint), then after 20 min of incubation either without (control) or with INK128 (300 nM) (**Fig. 1B, C**). INK128 (300 nM) did not cause any detectable cytotoxicity (WB using an antibody against caspase 3) but effectively decreased phosphorylation of S6 (serines 235/236) and Akt (serine 473), commonly used as indicators of mTORC1 and mTORC2 activity, respectively (**Fig. 1 D-G**). We counted the number of spines in which Rab11a-positive vesicles were present (**Fig. 1H, I**). Of these spines, an average of ∼66% contained mobile Rab11a that entered or exited the dendritic spine at some point over 1 min or moved from the dendritic spine head to the neck (**Fig. 1J, K**). The remaining spines contained only Rab11a, did not move, and resided primarily at the spine head (**Fig. 1J, K**; **Movies 1**, **2**). The thirty-minute preincubation step did not affect Rab11a dynamics. Twenty minutes of incubation with INK128 decreased the number of dendritic spines that contained mobile Rab11a by ∼30% (**Fig. 1J, K**) compared with the respective control. Altogether, these data indicated that mTOR inhibition, which is known to induce autophagy, resulted in the partial immobilization of Rab11a in dendritic spines.

**Fig. 1.**
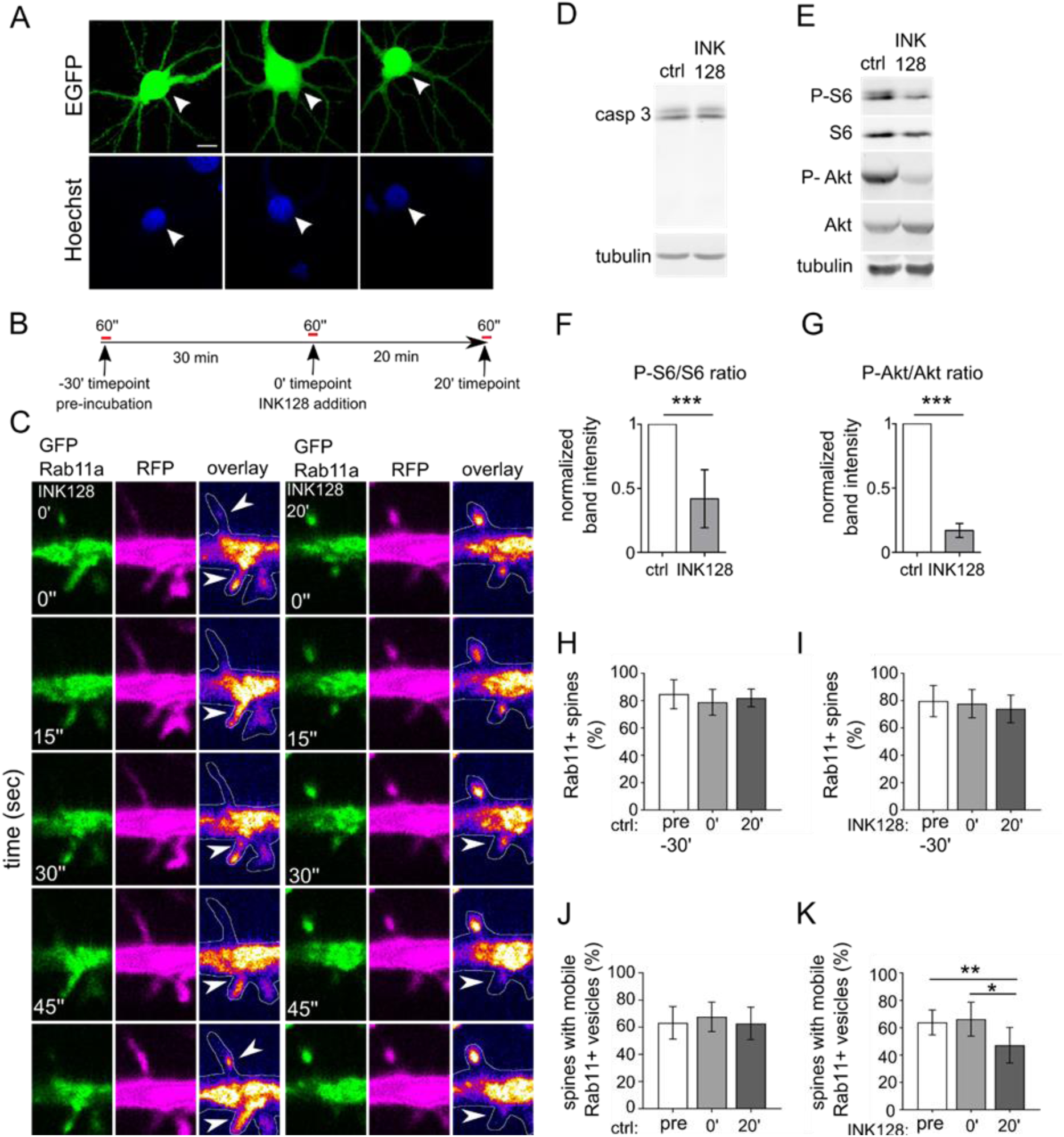
mTOR inhibition decreases Rab11 mobility in dendritic spines. (**A**) Representative images of neurons transfected, using Lipofectamine 2000, at ∼ 3 weeks in culture, with plasmid encoding EGFP (arrowheads) stained with Hoechst33258. The image shows healthy, non-shrunk nuclei. **(B)** Diagram of treatment and imaging for experiments in Fig 1. Neurons were recorded for 1 minute (preincubation recording; pre-30’), then incubated for 30 minutes to control potential phototoxicity or overheating in the microscope stage. Then, neurons were re-recorded (0’) and incubated for 20 minutes without (ctrl 20’) or with 300 nM mTOR inhibitor INK128 (INK128 20’) for the final recording. (**C**) Representative time-lapse images of dendrites of rat hippocampal neurons were cotransfected on DIV22 with plasmids encoding GFP-Rab11a and RFP and treated the next day with INK128 (300 nM). The figure shows Rab11a vesicles (green), RFP (magenta), an overlay with GFP-Rab11a (fire pseudocolor), and the outline based on the RFP channel (hand-drawn ROI). Arrows indicate dendritic spines with motile vesicles. Imaging occurred ∼17-24 h after transfection. Neurons were imaged 3 times, always for 1 min, at the baseline (not shown, in D, E; INK 128 pre-30’; see panels I, K), 30 min later just prior to treatment (INK128 0’) and after 20’ incubation with INK128 (INK128 20’). Control neurons (not shown in C) were imaged as described above without any treatment (ctrl pre-30’, ctrl 0’ and ctrl 20’, see H and J). Scale bar = 2.5 µm. (**D**) Western blot analysis of INK128 treatment effects on caspase 3 (casp 3) cleavage in cultured neurons. Primary hippocampal rat neurons (DIV21) were treated with INK128 (300 nM) for 20 minutes. Extracts were blotted against casp 3 with tubulin as a loading control. (**E-G**) Western blot analysis of mTORC1 and mTORC2 inhibition by INK128. The same extracts as in (D) were blotted against phosphorylated (Ser235/S236, P-S6) and total S6 and phosphorylated (Ser473, P-Akt) and total Akt to evaluate mTORC1 and mTORC2 suppression, respectively, with tubulin as a loading control. Data are presented as mean gray values for each phosphorylated protein normalized to total protein. Error bars show the standard deviation. *n* = 6 both for ctrl and INK128. \*\*\**p* ≤ 0.001 (one-sample *t*-test) **(H)** Percentage of Rab11a-positive spines in control neurons. (**I**) Percentage of Rab11a-positive spines in INK128-treated neurons. (**J**) Percentage of spines with mobile Rab11a in control neurons. (**K**) Percentage of spines with mobile Rab11a in INK128-treated neurons. The data are presented as a mean percentage of Rab11a positive spines normalized to all spines (H, I) or the mean percentage of Rab11a positive spines with mobile vesicles normalized to all Rab11a positive spines for all analyzed cells (J, K). Error bars show the standard deviation. *n* = 11 cells for treated and untreated neurons from four independent experiments. \**p* ≤ 0.05, \*\**p* ≤ 0.01 (repeated-measures ANOVA followed by Tukey’s *post hoc* test).

### Co-occurrence of Rab11 and Atg9A increases upon mTOR inhibition

The mobile pool of Rab11a in dendritic spines was previously assigned to recycling endosomes and the delivery of α-amino-3-hydroxy-5-methyl-4-isoxazolepropionic acid (AMPA)-type ionotropic glutamate receptors to postsynaptic sites (Esteves da Silva et al., 2015). The presence of stationary Rab11a, which is present in membranous structures, combined with the earlier findings on the potential contribution of Rab11 to autophagy, prompted us to investigate whether Rab11-positive vesicles participate in this process at dendritic spines upon mTOR inhibition. We evaluated changes in the co-occurrence of Rab11 with autophagy- and recycling endosome-associated markers in response to INK128 application. A frequently used autophagy marker is LC3, which is recruited to the forming APs and is present until mature APs reach lysosomes. LC3-positive APs are sparse at dendritic spines, which is speculated to result from their dynamic behavior and short lifespan (Shehata et al., 2012). Little is known about preautophagosomal structures and the steps upstream of phagophore formation in the dendrites and dendritic spines. Therefore, we used another marker, Atg9A, to visualize early endomembrane structures where autophagy can originate. Atg9A is a transmembrane protein that initiates autophagy and can migrate through recycling endosomes to autophagosome nucleation sites (Imai et al., 2016). It transiently colocalizes with autophagosomes in HEK293 cells (Orsi et al., 2012).

DIV21-25 rat hippocampal neurons were transfected with plasmid encoding EGFP to trace dendrites and spines of individual cells more accurately, treated with INK128 as described above, and fixed for immunofluorescence and super-resolution imaging. We performed double staining for Rab11 using three different markers. Besides Atg9A, as the marker of pre-autophagic vesicles, we stained neurons with Hook3 and syntaxin-12, which are associated with endocytic recycling. Hook3 is a cargo adaptor protein known to form a motile complex with dynein and dynactin in microtubular transport (Schroeder and Vale, 2016). Endosomal transport in dendritic spines mostly depends on actin filaments and actin-myosin interactions. However, microtubules were recently shown to dynamically enter and recede from dendritic spines in an activity-dependent manner, thus accelerating Rab11-positive vesicle transport (Esteves da Silva et al., 2015, Jaworski et al., 2009, Gu et al., 2008, Hu et al., 2008). Another marker associated with endocytic recycling is syntaxin-12 (also known as syntaxin 13), which binds to Rab11 through Glutamate receptor-interacting protein 1 (GRIP1)-associated protein 1 (GRASP-1) (Hoogenraad et al., 2010). Following antibody staining, we performed Airyscan super-resolution imaging to visualize the investigated proteins in dendritic spines. We have used the EGFP channel to separate the transfected neuron from the processes originating in other non-transfected neurons (**Fig. 2A**). Based on it, we isolated dendritic spines of all shapes with their spine neck and head, then performed an object-based colocalization analysis.

**Fig. 2.**
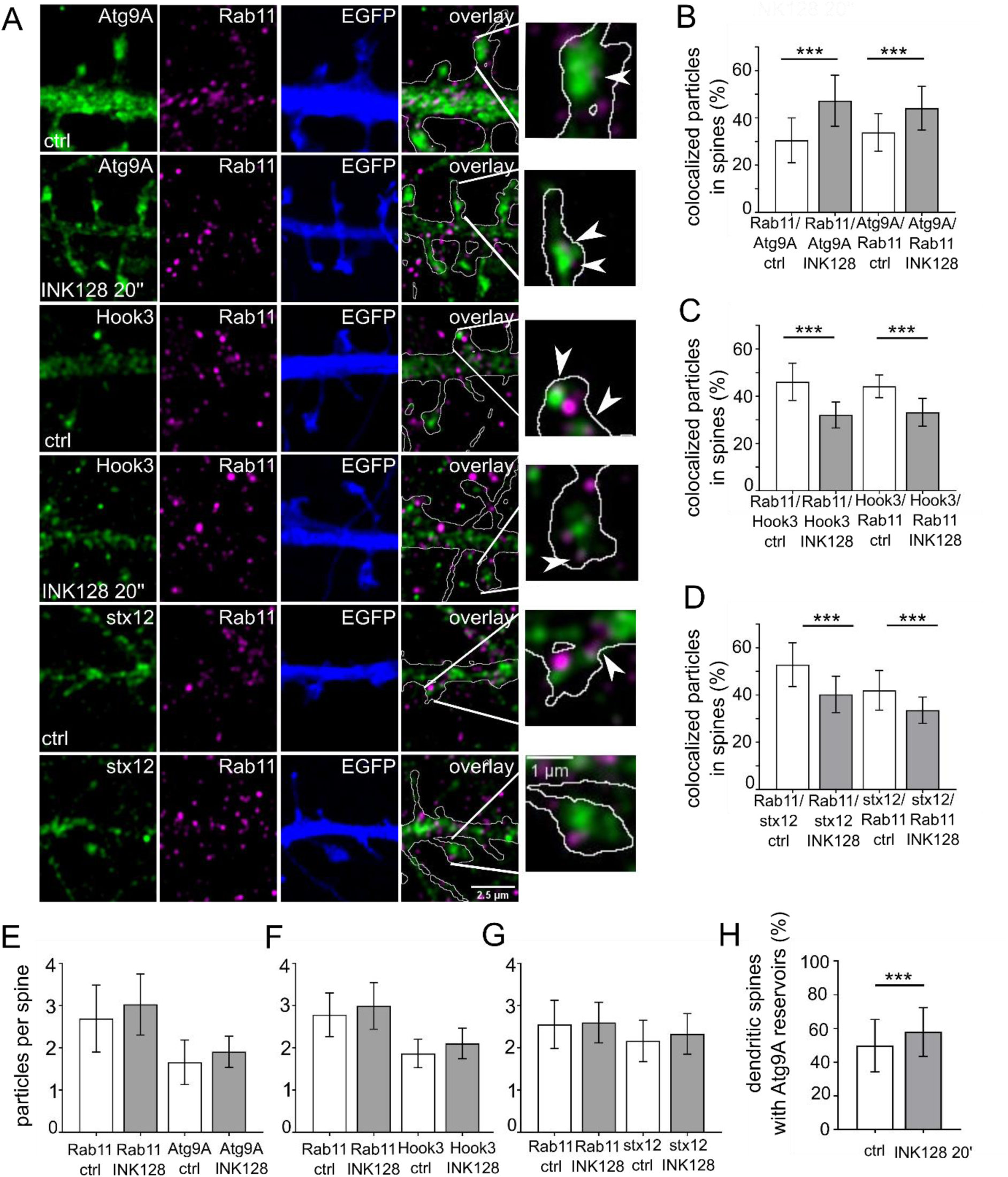
mTOR inhibition targets Rab11 to Atg9A-positive reservoirs. (**A**) Representative Airyscan images of dendrites of neurons that were transfected with a plasmid encoding EGFP (blue) on DIV22 to allow accurate dendritic spine tracing and immunostained the next day for native Rab11 (magenta) and Atg9A, Hook3, or syntaxin 12 (stx12) (green). The overlay channel shows Rab11 (magenta), Atg9A, Hook3 or stx12 (green), and the EGFP channel as an outline. The plasmid was expressed overnight. The neurons were incubated with INK128 (300 nM) for 20 min, fixed, and immunostained. Scale bar = 2.5 µm. The rightmost photomicrographs represent individual dendritic spines indicated in the overlay panels. Arrows indicate exemplary colocalized particles. Scale bar = 1 µm. (**B**) Colocalization of Rab11 with Atg9A in dendritic spines of neurons that were treated as in (A). (**C**) Colocalization of Rab11 with Hook3 in dendritic spines of neurons that were treated as in (A). (**D**) Colocalization of Rab11 with stx12 in dendritic spines of neurons that were treated as in (A). (**E**) Number of Rab11 and Atg9A particles per spine in neurons treated as in (A). (**F**) Number of Rab11 and Hook3 particles per spine in neurons treated as in (A). (**G**) Number of Rab11 and stx12 particles per spine in neurons treated as in (A). Data are presented as a mean percentage of colocalized particles normalized to all particles in each channel (B, C, D) or the number of total particles normalized to the number of dendritic spines (D, E, F) for all neurons. Error bars show the standard deviation. The number of cells for Atg9A immunofluorescence analysis: *n* = 32 (control) and 30 (INK128) neurons from four independent experiments. The number of cells for Hook3 immunofluorescence analysis: *n* = 28 for both variants from three independent experiments. The number of cells for stx12 immunofluorescence analysis: *n* = 24 for both variants from three independent experiments. \*\*\**p* ≤ 0.001 (one-way ANOVA followed by Sidak’s *post hoc* test). (**H**) Semi-quantitative estimate of the prevalence of Atg9A large vesicles (reservoirs) in the dendritic spines. Same neurons as in B were evaluated for the presence of Atg9A-positive structures > 0.150 µm^2^ in the spines. The result was normalized to the number of the dendritic spines and shown as a percentage. Data are presented as mean. *n* = 32 and 30 cells for control and INK128 variants from four independent experiments. \**p* < 0.05 (unpaired *t*-test).

Atg9A was common in dendritic spines, with either punctate staining or prominent structures primarily in heads of mushroom-shaped spines and occasionally at the base of the spine and in dendrites (**Fig. 2A**). At the baseline, over 40% of dendritic spines had large Atg9A structures (i.e., > 150 μm^2^), called Atg9 reservoirs. INK128 treatment slightly increased the fraction of such spines (**Fig. 2H**). The colocalization of Rab11 with Atg9A and *vice versa* increased from ∼30% to ∼45% after 20 min of INK128 treatment, while the average particle number per total number of spines did not significantly change (**Fig. 2B, E**). The reverse was true for Hook3, in which colocalization decreased from ∼45% to ∼32% (**Fig. 2C, F**). Rab11 colocalization with syntaxin-12 decreased by 12% upon INK128 treatment; this difference was smaller with syntaxin-12 *vs*. Rab11 colocalization (**Fig. 2D, G**). These results suggest that most syntaxin-12 co-occurs with Rab11 (a characteristic of recycling endosomes), while Rab11 changes its co-occurrence with syntaxin-12 upon INK128 treatment. Thus, the association between Rab11 and recycling endosomes was diminished. Altogether, the more significant co-occurrence of Rab11 and Atg9A and decrease in colocalization with the mobility marker Hook3 and recycling endosome marker syntaxin-12 suggest that Rab11 partly shifts to compartments that contribute to early autophagy upon mTOR inhibition. Overall, these results suggest that Rab11 plays a role in initiating autophagy in dendritic spines.

### Rab11a activity is necessary for Atg9A maintenance at dendritic spines

Rab11 and Atg9A co-occurred at dendritic spines, and their colocalization changed upon mTOR inhibition. We also observed that large Atg9A-positive structures are widespread at the dendritic spines. We next sought to confirm that the interaction between Rab11 and Atg9A has functional consequences in dendritic spines. The expression of a dominant-negative (DN) variant of Rab11a (S25N) blocks the activity of endogenous Rab11a, halts endocytic recycling, and changes the morphology of dendritic spines (Esteves da Silva et al., 2015). In mouse embryonic fibroblasts, Atg9A moved through recycling endosomes that delivered it to preautophagosomal sites (Imai et al., 2016), suggesting that Rab11 might be needed for Atg9A to reach its destination. Therefore, we investigated whether Rab11a activity is necessary to maintain Atg9A at dendritic spines. We transfected DIV21-25 rat embryonic hippocampal neurons with plasmids encoding iRFP702 to fill the dendrites using Scarlet-I-Atg9A, and one of the three variants of GFP-Rab11a (wildtype [WT], DN Rab11a variant S25N (Hunyady et al., 2002), or constitutively active [CA] Rab11a variant S20V) (Pasqualato et al., 2004). The next day, we analyzed live neurons’ Scarlet-I-Atg9A distribution and colocalization with GFP-Rab11a variants (by object-based colocalization analysis). Scarlet-I-Atg9A was present in dendritic spines of GFP-Rab11a WT and GFP-Rab11a CA transfected neurons, where it colocalized with GFP-Rab11 were no significant differences between analyzed variants (**Fig. 3A-C**; **Movies 3, 4**). In Rab11a DN overexpressing neurons, however, Scarlet-I-Atg9A presence in dendritic spines decreased from ∼0.9 to 0.45 particles per dendritic spine number (i.e., all dendritic spines in focus in the field of view) compared with Rab11a WT. Also, Scarlet-I-Atg9A/GFP-Rab11a colocalization in dendritic spines greatly diminished in Rab11a DN transfected cells (from ∼80% to ∼30%; **Fig. 3A-C**). Of note, GFP-Rab11a DN was present and diffusely distributed in all dendritic spines of transfected neurons (**Fig. 3A**); therefore, the results of colocalization, in this case, depend only on the dendritic distribution of Atg9A and not its co-occurrence with the recycling endosomes. Altogether, these data showed that Rab11a activity was needed for Atg9A maintenance at dendritic spines.

**Fig. 3.**
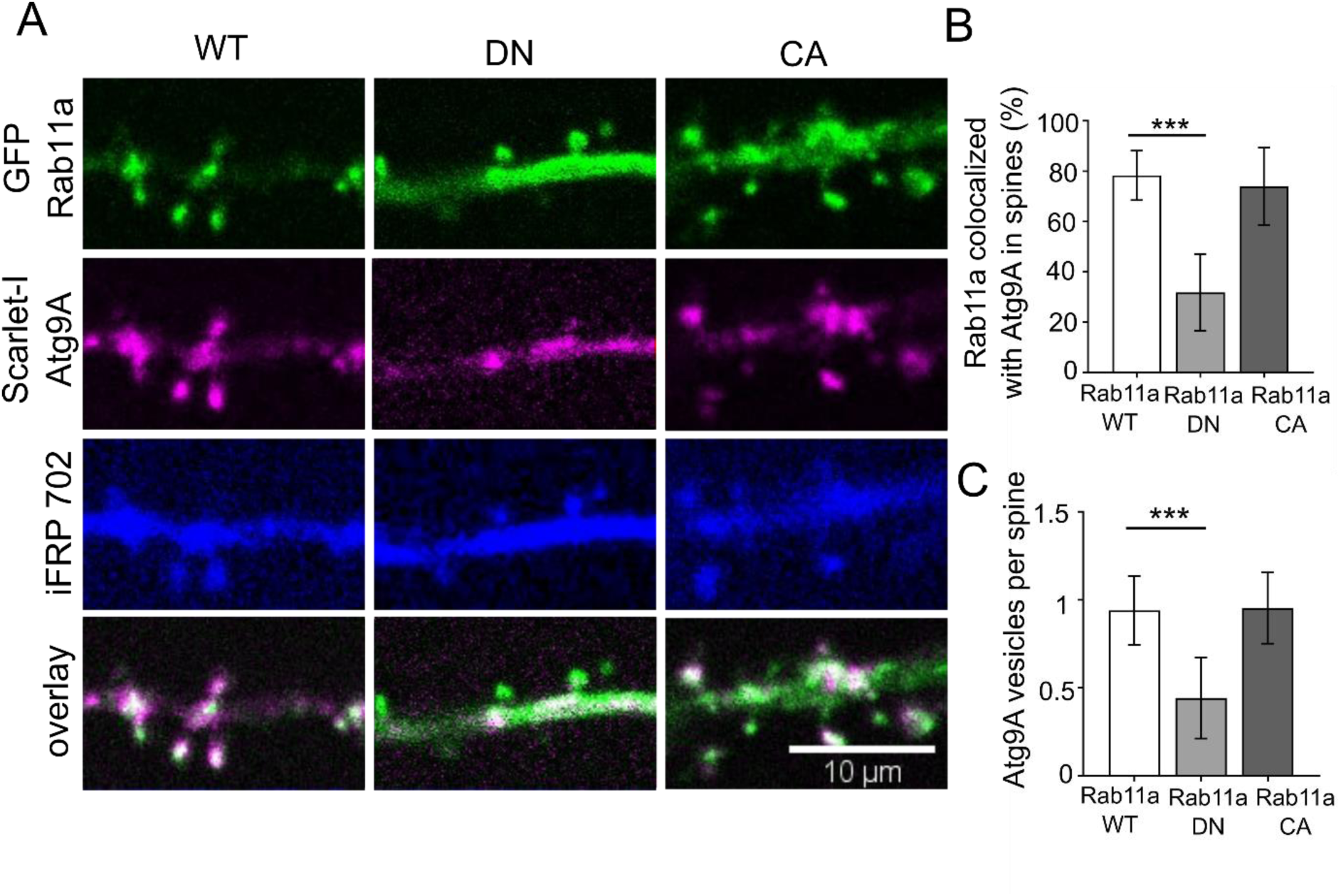
Rab11a GTPase activity is necessary for the presence of Atg9A at dendritic spines. (**A**) Representative images of DIV22 neurons that were cotransfected with plasmids encoding GFP-Rab11a, dominant-negative (DN) Rab11a, or constitutively active (CA) Rab11a (green) together with Scarlet-I-Atg9A (magenta) and iRFP702 (blue). Imaging occurred 17-24 h after transfection. Scale bar = 10 µm. (**B**) Colocalization rate (percentage of colocalized vesicles) of transfected Rab11 variant and Atg9a at dendritic spines. (**C**) The number of Atg9a particles per spine of neurons transfected with the indicated Rab11 variants. Data are presented as the mean percentage of colocalized Rab11a particles normalized to all particles (B) or the number of Atg9A particles normalized to the number of spines for all cells (C). Error bars show the standard deviation. Number of analyzed cells: *n* = 30 for Rab11a WT, *n* = 30 for DN Rab11, and *n* = 27 for CA Rab11a from three independent experiments. \*\*\**p* ≤ 0.001 (one-way ANOVA followed by Sidak’s *post hoc* test).

### Rab11 co-precipitates with Atg9A in mouse synaptoneurosomes

Since we demonstrated that Rab11 and Atg9A partially colocalize, we sought to investigate if these two proteins occur in the same protein complex. To this end, we additionally performed Rab11 co-immunoprecipitation with an anti-Rab11 antibody. Since Atg9A is not indicative of ongoing autophagy and only provides the capacity to begin the autophagy, we were also interested in an established autophagy marker, namely LC3. Therefore, we sought to investigate whether LC3B, an isoform of LC3 that is widely expressed in the brain, is co-precipitated by Rab11, which would confirm the study by Puri et al. in HeLa cells (2018). As starting material, we used synaptoneurosomes, a biochemical model of the synapse that previously worked in our hands (Janusz et al., 2013). Other researchers reported the participation of autophagy in cLTD. Considering the results of our earlier experiments with INK128, we also investigated whether the degree of Atg9A co-immunoprecipitation with Rab11 in synaptoneurosomes changes upon NMDA stimulation that in other works evoked cLTD (Lee et al., 1998, Li et al., 2004) or mTOR suppression.

Synaptoneurosomes that were derived from cortices and hippocampi of adult mice (Janusz et al., 2013) were treated with INK128 (600 nM), NMDA (50 μM), or a combination of both drugs. Treatment with NMDA was used to mimic cLTD induction in cultured neurons. Instead of washing out the NMDA after 5 min of stimulation, synaptoneurosomes were diluted 5x to decrease the concentration of NMDA below levels shown to evoke cLTD (Lee et al., 1998) and then incubated for 25 minutes. Indeed, this kind of treatment decreased the level of serine 845 phosphorylated GluA1 (P-GluA1), a subunit of AMPA receptors (**Fig. 4A, B**), which is considered the biochemical hallmark of cLTD (Kameyama et al., 1998, Lee et al., 1998). **Fig. 4C, D** shows that both Atg9A and LC3B co-immunoprecipitated with Rab11. INK128 alone had no apparent influence on the Rab11/Atg9A interaction in synaptoneurosomes likely due to its weak effect on P-S6 (Ser235/236) in our synaptoneurosome preparations (**Fig. 4E, F**). NMDA treatment tended to increase Rab11-Atg9A immunoprecipitation, but a significant difference was achieved only when combined with INK128 (**Fig. 4D**), suggesting that Rab11 associates more efficiently with Atg9A at the synapse upon cLTD induction. However, no such effect was observed with LC3B, a classic marker of autophagy (**Fig. 4G, H**). These findings prompted us to evaluate the emergence and dynamics of autophagosomes in dendritic spines upon conditions known to induce cLTD in live hippocampal neurons.

**Fig. 4.**
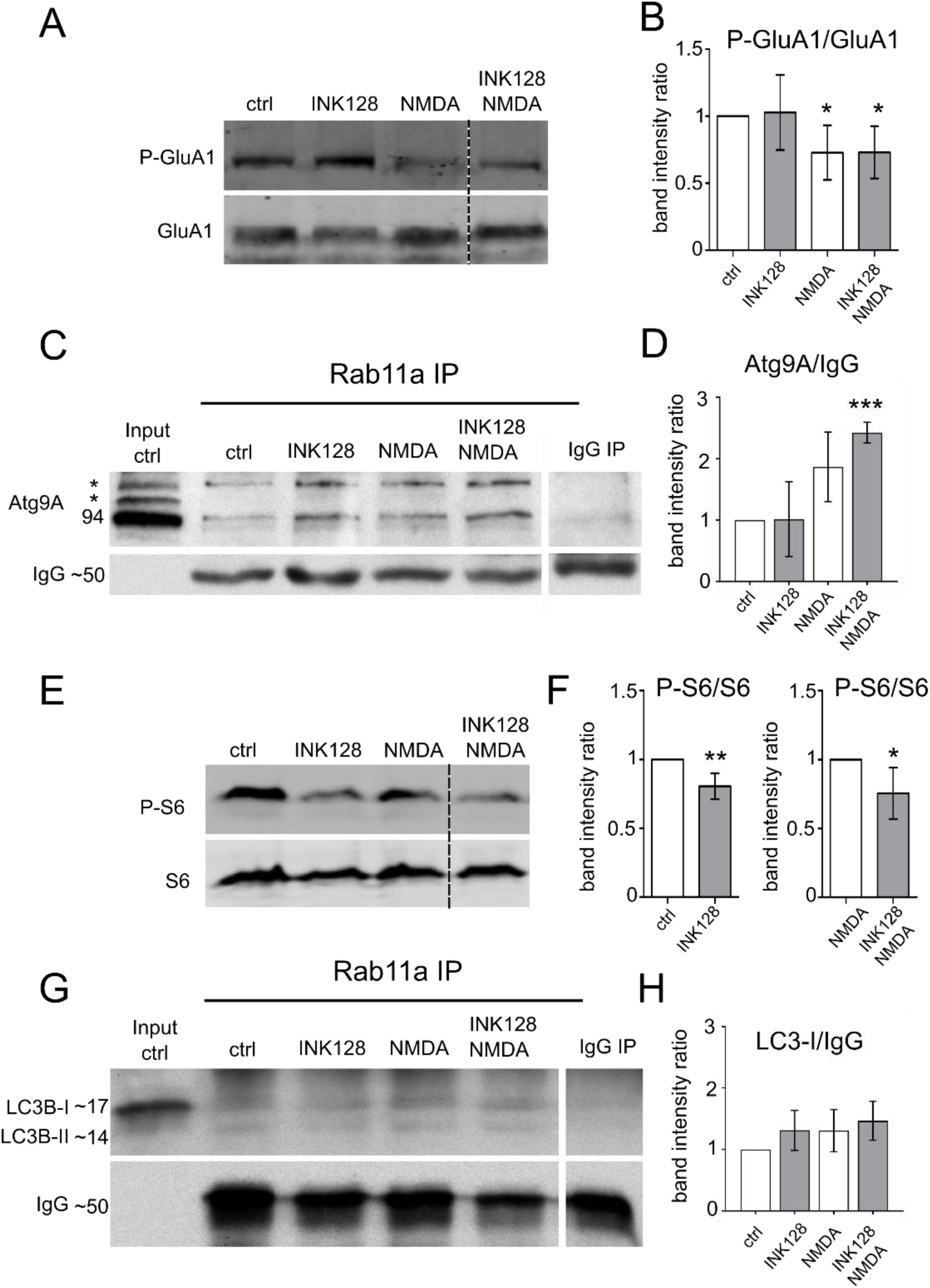
Rab11a interacts with Atg9A in mouse synaptoneurosomes. (**A, B**) Western blot analysis of NMDA-cLTD induction in synaptoneurosomal preparations. Synaptoneurosomes were prepared as for Figure 4, and levels of protein extracts were blotted against phosphorylated (Ser845; P-GluA1) and total GluA1 to evaluate GluA1 dephosphorylation considered a marker for cLTD. Dashed lines indicate membrane image cuts. Data are presented as mean gray values for each phosphorylated protein normalized to total protein. Error bars show the standard deviation. *n* = 5 for all samples. \**p* < 0.05, \*\**p* < 0.01 (one-sample *t*-test). (**C**) Western blot analysis of Atg9A co-immunoprecipitation with Rab11a from mouse synaptoneurosomes. Samples were first subjected to the treatments and then co-immunoprecipitated with anti-Rab11a antibody (see Methods). The blot shows the control immunoprecipitation sample following the input fraction (precleared control synaptoneurosome extract). The samples were then treated with INK128 (600 nM) alone, NMDA (50 µM) alone, or NMDA+INK128. The bands marked as * corresponds to posttranslational modifications of Atg9A. (**D**) Atg9A signal intensity ratios of the respective samples. The lower band (Atg9A without posttranslational modifications) was used for quantification. Mean gray values were normalized to IgG as a loading control and then to the control sample. Data are presented as a mean from all biological replicates. Error bars show the standard deviation. *n* = 4. \*\*\**p* ≤ 0.001 (one-sample *t-*test). (**E, F**) Western blot analysis of mTORC1 inhibition by INK128. Synaptoneurosome extracts were blotted against phosphorylated P-S6 (Ser235/236; C, D) and S6 to evaluate mTORC1 complex suppression. Dashed lines show membrane image cuts. Data are presented as mean gray values for each phosphorylated protein normalized to total protein. Error bars show the standard deviation. *n* = 5 for all samples. \**p* < 0.05, \*\**p* < 0.01 (one-sample *t*-test). (**G**) Western blot analysis of LC3B co-immunoprecipitation with Rab11a from mouse synaptoneurosomes prepared and treated as in A. **(H)** LC3B signal intensity ratios, calculated as above. The band corresponding to LC3-I was used for quantitation. Data are presented as a mean from all biological replicates. Error bars show the standard deviation. *n* = 3.

### Autophagy is enabled in dendritic spines of rat hippocampal neurons by simultaneous mTOR inhibition and NMDA treatment

We have yet to determine whether the same treatments that increase the Atg9A and Rab11 interaction induce autophagosome formation in dendritic spines. Therefore, we experimented with live primary neuronal hippocampal cultures. We counted LC3 puncta as a proxy of autophagosome formation at the dendritic spines as they appeared in real-time after stimulation, without applying any treatments that block autophagosome progression to downstream structures. We transfected cells with a pMX-IP-EGFP-LC3 plasmid, taking advantage of the retroviral long terminal repeat as a weak promoter: therefore, having less potential to overexpress LC3 to the degree at which it forms aggregates. We used the EGFP-tagged version of LC3 despite the recent report of high background in dendrites (Kulkarni et al., 2021) since we have not observed this unspecific staining in spines. We performed confocal spinning-disc imaging and treatments (i.e., application of INK128 or/and NMDA, proven previously to induce cLTD-like changes in neurons cultured *in vitro*, e.g., AMPA receptors internalization (Hsin et al., 2010, Li et al., 2010, Lee et al., 2002), the next day after transfection. Our treatments triggered neither cell death nor increased autophagy at the whole-cell level (**Fig. 5A**), thus suggesting that the newly formed APs observed in the dendrites and dendritic spines approximated a physiological reaction. For quantitative measurement of the autophagic response at the postsynapse, new APs were counted as one event, regardless of whether they remained at the spine, withdrew to the dendrite, or shuttled for several minutes between the spine and dendrite (e.g., **Movies 5-8**). New autophagosomes appeared at dendritic spines as soon as 10-15 min upon NMDA application and washout in the presence of INK128 (**Fig. 5B, G**). New autophagosomes were less common in the other stimulation variants, i.e., NMDA or INK128 alone (**Fig. 5G**). A biochemical approximation of cLTD induction by NMDA treatment was demonstrated by decreased levels of phosphorylated GluA1 (P-S845) (Kameyama et al., 1998, Lee et al., 1998), while INK128 resulted in a substantial decrease in mTOR activity (**Fig. 5C-F**). Our observations revealed that the neurons reacted differently to treatments depending on whether they already had multiple AP puncta at the dendritic spines before the stimulation. To tease out this difference, we subdivided neurons into two groups: neurons with low local autophagy at the start of the experiment (i.e., ≤ 6 dendritic spines that contained autophagosomes) and neurons with high local autophagy at the start of the experiment (i.e., > 6 dendritic spines that contained autophagosomes; **Fig. 5H, I**). Altogether, our results revealed that cLTD-like NMDA stimulation induced the emergence of LC3 autophagosome-like puncta at dendritic spines shortly after application and that a large subpopulation of them reacted when enabling conditions (i.e., low mTOR activity) were met.

**Fig. 5.**
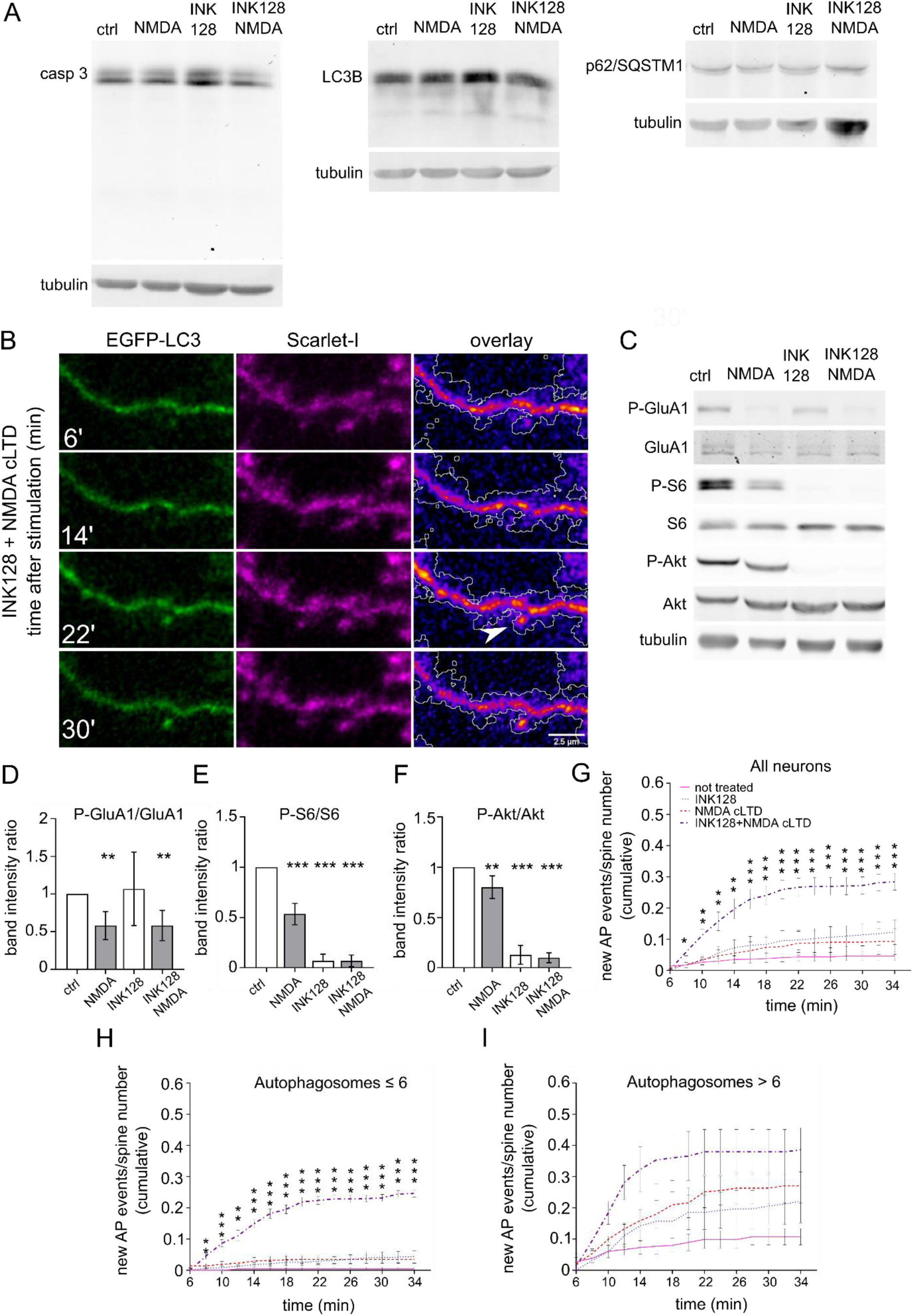
Simultaneous mTOR inhibition and NMDA-dependent cLTD induces the emergence of autophagosomes at dendritic spines in rat hippocampal cultured neurons. (**A**) Western blot analysis of levels of caspase 3 (casp 3), LC3B, and p62 in primary hippocampal rat neurons (DIV21) treated with INK128 (300 nM) or DMSO for 15 min and then NMDA (50 µM) for 3 minutes followed by the medium changed, and incubation with DMSO or INK128 (300 nM) for next 35 min. Tubulin is shown as a loading control. (**B**) Representative images of a dendrite of a rat hippocampal neuron cotransfected with plasmids encoding EGFP-LC3 and Scarlet-I on DIV22 were treated the next day with INK128 (300 nM) and NMDA (50 µM). The arrow points to a new autophagosome emerging upon treatment. Scale bar = 2.5 µm. (**C-F**) Western blot analysis of NMDA-cLTD induction and mTORC1 and mTORC2 inhibition by INK128. Extracts treated as in (A) were blotted against phosphorylated (Ser845; P-GluA1) and total GluA1 to evaluate GluA1 dephosphorylation considered a marker for cLTD, P-S6 (Ser235/236), and S6 to evaluate mTORC1 complex suppression and P-Akt (Ser473) and Akt to evaluate mTORC2 suppression. Tubulin level serves as a loading control. Data are presented as mean gray values for each phosphorylated protein normalized to total protein. Error bars show the standard deviation. *n* = 7 for all samples. ** *p* ≤ 0.01, *** *p* ≤ 0,001 (one-sample *t*-test). **(G)** Neurons that were transfected as in (B) were either untreated or pretreated with INK128 (300 nM) for 15 min and then treated with NMDA for 3 min. The medium was then changed, and the neurons were incubated with or without INK128 (300 nM). Cumulative analysis of EGFP-LC3 vesicles showed that they appeared at dendritic spines at 2 min intervals, starting 6 min after NMDA application (shown as the total number of events normalized to the number of visible spines). Data are presented as a mean of cumulative events normalized to the number of spines for all time points and cells. Error bars show the standard error of the mean. *n* = 14 control cells. *n* = 12 INK128-treated cells. *n* = 10 NMDA-treated cells. *n* = 11 INK128+NMDA-treated cells. \**p* ≤ 0.05, \*\**p* ≤ 0.01, \*\*\**p* ≤ 0.001 (two-way ANOVA followed by Dunnett’s *post hoc* test). (**H, I**) Cumulative analysis of EGFP-LC3 vesicles of the same neurons as in (G), split into two groups with initial autophagosome number ≤ 6 (H) or > 6 (I). Data are presented as a mean of cumulative events normalized to the number of spines for all time points and cells. Error bars show the standard error of the mean. In (H): *n* = 11 for control cells, *n* = 7 for INK128-treated cells, *n* = 7 for NMDA-treated cells, *n* = 8 for INK128+NMDA-treated cells. In (I): *n* = 3 for control cells, *n* = 5 for INK128-treated cells, *n* = 3 for NMDA-treated cells, *n* = 3 for INK128+NMDA treated cells. \**p* ≤ 0.05, \*\**p* ≤ 0.01, \*\*\**p* ≤ 0.001 (two-way ANOVA followed by Dunnett’s *post hoc* test).

### EGFP-LC3B-negative spines are more dynamic in response to NMDA treatment

In the last series of experiments, we learned the fate of dendritic spines that formed autophagosomes in response to INK128 and NMDA. To this end, we experimented in an analogous way to that above, except that the imaging was prolonged by 10 minutes (i.e., to 46 min after stimulation) to observe the change in the shape of the dendritic spines (**Fig. 6A**). Analysis of dendritic spine length showed that dendritic spines in which EGFP-LC3 was absent in response to INK128 and NMDA administration changed their shape significantly more often than EGFP-LC3-positive ones (**Fig. 6B**). Therefore, the appearance of autophagosomes does not immediately lead to spine pruning; on the contrary, it correlates with their increased structural stability.

**Figure 6.**
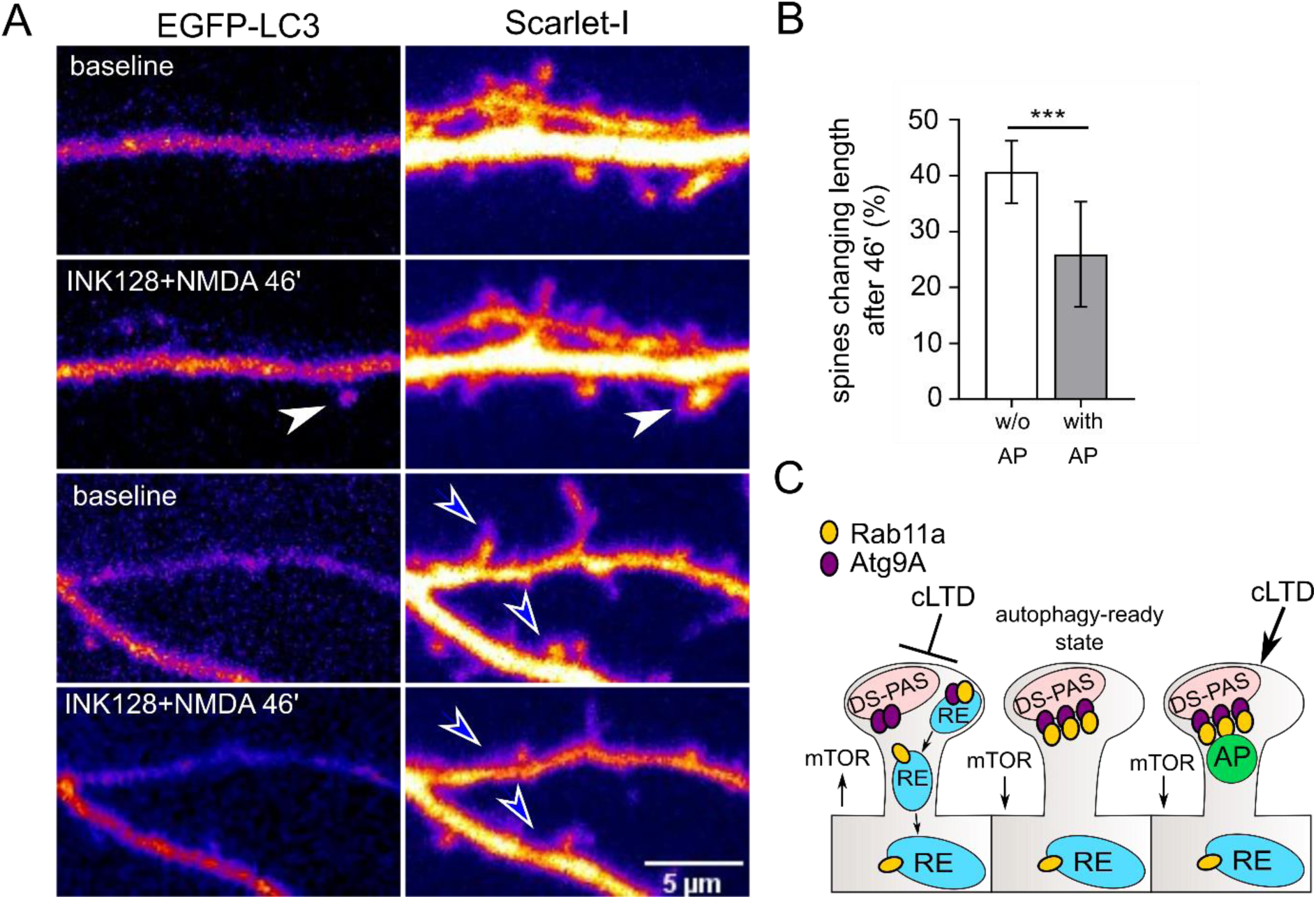
Dendritic spines that contain EGFP-LC3-positive vesicles after NMDA cLTD are less susceptible to shape change. (**A**). Representative images of dendrites of a rat hippocampal neuron cotransfected on DIV22 with plasmids encoding GFP-LC3 and Scarlet-I were treated with INK128 and NMDA the next day. Scale bar = 5 µm. Before treatment, neurons were recorded for 10 minutes (baseline). Then neurons were pretreated with INK128 (300 nM) for 15 minutes and next stimulated with NMDA (50 µM) for 3 min (INK128+NMDA 46’). Afterward, media was replaced with conditioned media containing INK128. Neurons were recorded every 2 min for 46 minutes, starting 6 min after NMDA application. Dendritic spines were evaluated for the presence of LC3-positive autophagosomes (APs) throughout the movie. White arrows show examples of dendritic spines where an AP appeared. Blue arrowheads show examples of dendritic spines without AP that underwent pruning. (**B**) Percentage of spines that underwent over 25% change in length compared to the baseline, evaluated for the emergence of APs through the time-lapse. Data are presented as a mean percentage of spines that underwent shape change for all cells. Error bars show the standard deviation. *n* = 9 cells from 4 independent experiments. \*\*\**p* ≤ 0.001 (unpaired *t*-test). (**C**) Proposed mechanism of two-step autophagy regulation by Rab11. At baseline mTOR activity, Rab11-positive recycling endosomes deliver Atg9A to its dendritic spine reservoir (DS-PAS). Upon mTOR inhibition, endocytic recycling is diminished, and Rab11 is more associated with DS-PAS, showing an autophagy-ready state. Upon cLTD, autophagy machinery assembles at the Rab11 platform, and DS-PAS supplies the membrane to form an AP.

## Discussion

The present study showed that Rab11 GTPase co-occurs with Atg9A at dendritic spines, and this co-occurrence increases upon mTOR suppression. We identified an Atg9A reservoir at dendritic spines, and we propose that it serves as a site of nucleation for autophagosomes. We argue that Rab11 regulates the maintenance of this reservoir and that the Rab11/Atg9A interaction could be a prerequisite for subsequent autophagy at dendritic spines induced by neuronal activity. We also propose that autophagosome formation at the postsynapse is a common physiological process that occurs after cLTD and is enabled by low mTOR kinase activity.

The entrance of Rab11 into dendritic spines in the context of neuroplasticity, namely chemical LTP, was described previously; Rab11-positive recycling endosomes participated in returning AMPA receptors to the postsynapse (Esteves da Silva et al., 2015). We found that the suppression of mTOR activity diminished the movement of Rab11-positive vesicles, but this did not influence their presence at the postsynapse in the short term (i.e., up to 40 min). We also showed that Rab11 colocalized with Atg9A (**Fig. 2**), similar to non-neuronal cells, and confirmed their biochemical association by co-immunoprecipitation (**Fig. 4**). This interplay between Rab11 and Atg9A depends on both mTOR activity and stimulation. Rab11 and Atg9A colocalization increased within 20 min upon mTOR inhibition in neuronal cultures, while colocalization with other proteins related to endocytic transport and recycling decreased (**Fig. 2**). In synaptoneurosomes, a preparation considered a biochemical model of the synapse, the increase in the Rab11/Atg9A association depended on NMDA application (**Fig. 4**). These findings suggested that Rab11 partially switched from recycling endosomes to immobilized compartments, which are potentially autophagosome nucleation sites, and that stationary Rab11-positive vesicles could participate in autophagy.

The hypothesis that Rab11 participates in autophagy was tested in several studies of non-neuronal cells. Rab11 was required for autophagy in HEK293A cells and contributed to the early steps of autophagosome formation via Atg9A and serine/threonine-protein kinase Ulk1 (Longatti et al. 2012). Recycling endosomes have a role in Atg9A and Atg16L compartment fusion, leading to autophagy (Puri et al. 2013). Also, Atg9A-containing membranes are generated at Rab11 reservoirs in HeLa cells and mouse fibroblasts (Takahashi et al., 2016). Finally, the Rab11a compartment may serve as a platform to assemble other proteins for autophagy machinery (Puri et al., 2018). These studies showed that Rab11 is crucial for autophagy, notably autophagy initiation in mammalian cells.

Rab11 might play a dual role in initiating autophagy in dendritic spines as a vehicle for transporting autophagy proteins and as an autophagy platform. We showed that the presence of Atg9A reservoirs at dendritic spines in the long term (i.e., after overnight transfection) depends on Rab11 GTPase activity, suggesting that the postsynaptic Atg9A pool must be regularly replenished by recycling endosomes. Our results are consistent with other cell lines’ findings (Longatti et al., 2012, Puri et al., 2013, Imai et al., 2016).

One issue is whether the Rab11/Atg9A assembly increase at dendritic spines is equivalent to autophagosome formation. We applied the same treatments, mTOR suppression with INK128, and NMDA cLTD induction individually and together in a live imaging experiment with EGFP-LC3 transfected neurons to resolve this question. We showed that Rab11/Atg9A colocalization increases upon mTOR suppression in cultured neurons. However, INK128 treatment alone did not cause the formation of nascent autophagosomes in the dendritic spines of neurons with a low baseline number of autophagosomes at the spines. Thus, we can infer that increased colocalization of Rab11/Atg9A is not equivalent to autophagy. NMDA-cLTD induction alone also did not cause the emergence of autophagosomes in the dendritic spines of neurons with a low baseline number of autophagosomes. However, simultaneous mTOR suppression and NMDA-induced cLTD induced the emergence of autophagosomes at the dendritic spines in the case of neurons with low baseline AP count at the spines. When baseline autophagy in cells was already at a higher level, new autophagosomes arrived at spines regardless of which treatment was applied. This observation may explain the differences between our work and research performed by Shehata et al. (2012) or Kallergi et al. (2022), in which cLTD was sufficient for increased autophagosome presence in the dendrites and dendritic spines. While these studies were based on fixed neurons, we were the first to observe the same cells over time after the stimulation, revealing subpopulations of high and low-autophagosome neurons. Because the neurons with high basal AP count reacted strongly to any treatment, it is easy to see how they could obscure the population with low basal AP count in the other studies. Furthermore, our division of neurons into those two groups is arbitrary, so the difference between the groups is likely to be gradual. Nevertheless, our experiments show that basal mTOR activity is an essential factor that regulates neuronal response to the NMDA in the case of autophagy.

In the present study, we were also able to observe the dynamics and location of the nascent autophagosomes. Most new autophagosomes at the dendritic spines emerged within 10-18 min after NMDA treatment. Using 2-min intervals to prevent bleaching prevented us from discerning which autophagosomes originated in situ and which ones arrived there. However, the synaptoneurosome co-immunoprecipitation experiment supported the hypothesis that autophagosomes can arise at the postsynapse because LC3B is present within the Rab11 complex at dendritic spines. Only local proteins participate in the Rab11/Atg9A/LC3B complex in synaptoneurosome preparations as they cannot arrive from other cellular compartments. This does not exclude the formation of the AP in the dendritic shaft or other regions of the dendrite, as indicated by our colleagues (Shehata et al., 2012, Kallergi et al., 2022). Although we have not focused on Atg9A reservoirs in the dendrites, we speculate that neurons are likely to possess protein processing compartments located in their other parts, including shaft and branching points.

Overall, our results suggest that dendritic spines have the potential to perform cLTD-dependent autophagy on demand. We propose that this process is regulated by Rab11 and mediated by the Atg9A reservoir that we tentatively call the dendritic spine pre-autophagosomal structure (**Fig. 6E**). Unknown is whether this compartment corresponds to the spine apparatus or other endomembranes. The lower basal levels of mTOR activity may be a prerequisite but are not the cause of physiological autophagy at postsynaptic sites. We propose a model in which the initial steps that lead to mTOR-dependent autophagosome machinery assembly are left on hold in dendritic spines and are selectively resumed upon neuronal stimulation. Our results suggest that autophagosomes do not lead to immediate spine retraction or elongation into filopodia, as hypothesized by others (Kallergi et al., 2022). We speculate that dendritic spines where autophagosomes emerge are recruited from the pool of well-developed, mature spines that contain the necessary machinery to recruit autophagy proteins, and these are not prone to the rapid changes in shape. Autophagosome function in NMDA-evoked cLTD might be more subtle than degrading the whole dendritic spine. Dendritic autophagy removes AMPA receptors, an essential step for LTD (Shehata et al., 2012, Kallergi et al., 2022).

Moreover, recent studies reported that cLTD-induced autophagosomes contain many more proteins than AMPAR subunits, including elements vital for neuronal plasticity and spine structure, e.g., PSD-95 (Compans et al., 2021, Kallergi et al., 2022). These findings suggest that cLTD-triggered spine content remodeling may be essential to morphological change of initially stable spines. Kulkarni et al. (2021) showed that dendritic autophagy is also triggered rapidly by neuronal stimulation known to increase synaptic strength and spine growth. Thus, questions arise if autophagy occurs in proximity to “plastic” less stable spines under such conditions and how autophagosomal content is different from that revealed recently in the case of cLTD (Kallergi et al., 2022). These questions, however, remain beyond the scope of this study.

## Author Contributions

AJ-K and JJ designed the experiments. AJ-K, AB, AT, MB, MU, JM, JZ, and KK performed and analyzed the experiments. AJ-K and JJ discussed the results. AJ-K and JJ prepared the manuscript. JJ secured funding.

## Acknowledgments

We thank Bartosz Tarkowski for help with the SLIC primer design, Alina Zielinska and Kinga Kuchcinska for technical support, and Michael Arends for proofreading the manuscript. We also thank Dr. Magdalena Dziembowska and Dr. Gary Bassell for their comments on the manuscript. The Polish National Science Centre Opus supported this work with grants nos. 2012/07/E/NZ3/00503 and 2017/27/B/NZ3/01358 to JJ. AB was partly supported by a Polish National Science Centre Opus grant no. 2016/21/B/NZ3/03639 to JJ. AT was partly financed by a Polish National Science Centre Opus grant no. 2017/25/N/NZ3/01280. JJ, MU, and JZ are partly or wholly financed by the TEAM grant from the Foundation for Polish Science (POIR.04.04.00-00-5CBE/17-00).

## Abbreviations

ANOVA: analysis of variance
AMPA: α-amino-3-hydroxy-5-methyl-4-isoxazolepropionic acid
AP: autophagosomes
Atg: autophagy-related protein
CA: constitutively active
cLTD: chemical long-term depression
DIV: day *in vitro*
DN: dominant-negative
EGFP: enhanced green fluorescent protein
GTPase: guanosine triphosphatase
HEK293A: human embryonic kidney 293A
HRP: horseradish peroxidase
LC3: microtubule-associated protein 1A/1B-light chain 3
mTOR: mammalian/mechanistic target of rapamycin
mTORC1: mTOR complex 1
NMDA: N-methyl-D-aspartate
PBS: phosphate-buffered saline
Rab11: Ras-related protein Rab-11A
RFP: red fluorescent protein
ROI: region of interest
SDS-PAGE: sodium dodecyl sulfate-polyacrylamide gel electrophoresis
WB: Western blot

## Multimedia

**Movie 1.**
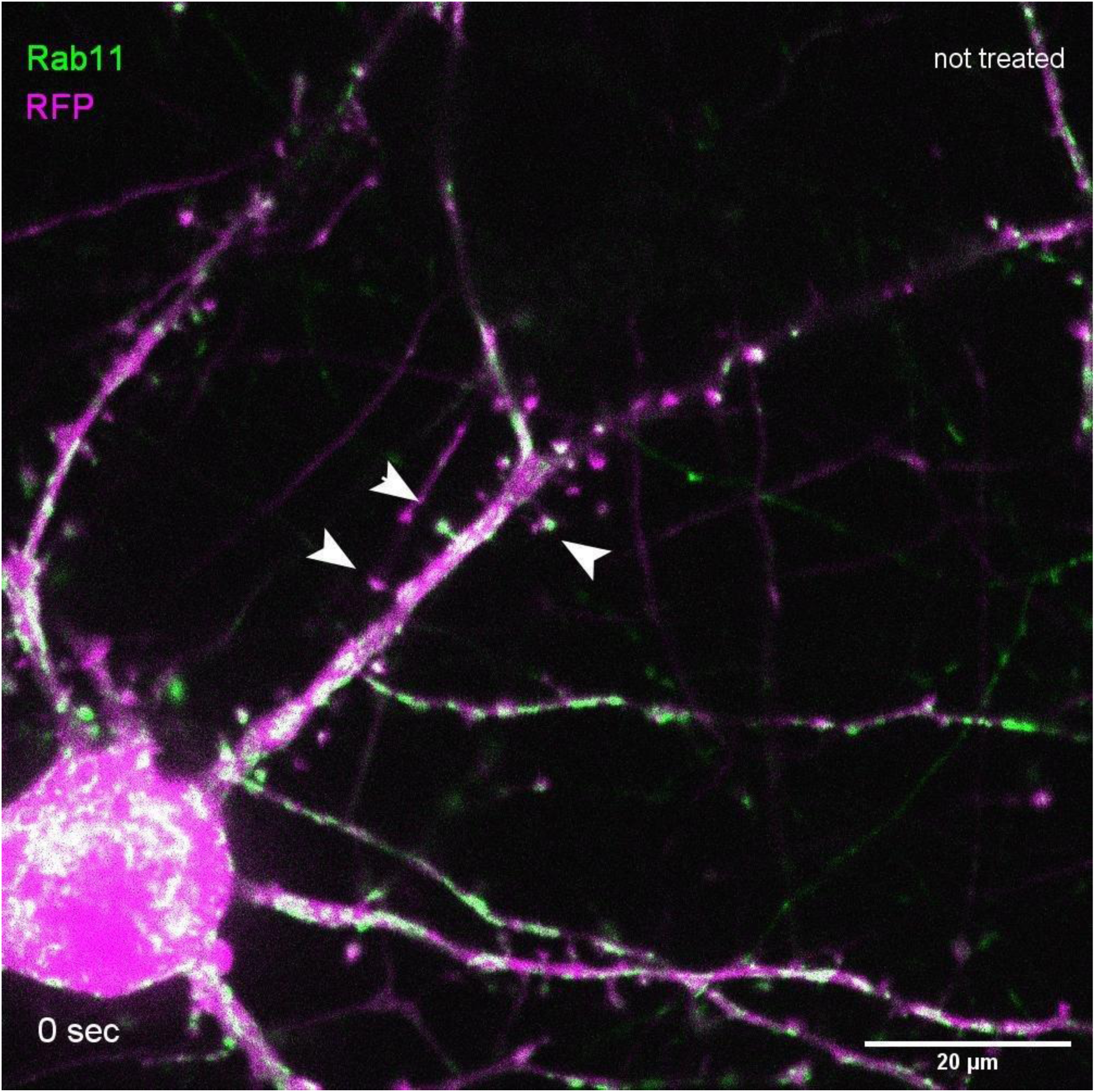
Hippocampal neuron transfected with plasmids encoding GFP-Rab11a (green) and RFP (magenta) over 1 min. Arrows point to examples of spines where mobile vesicles are present.

**Movie 2.**
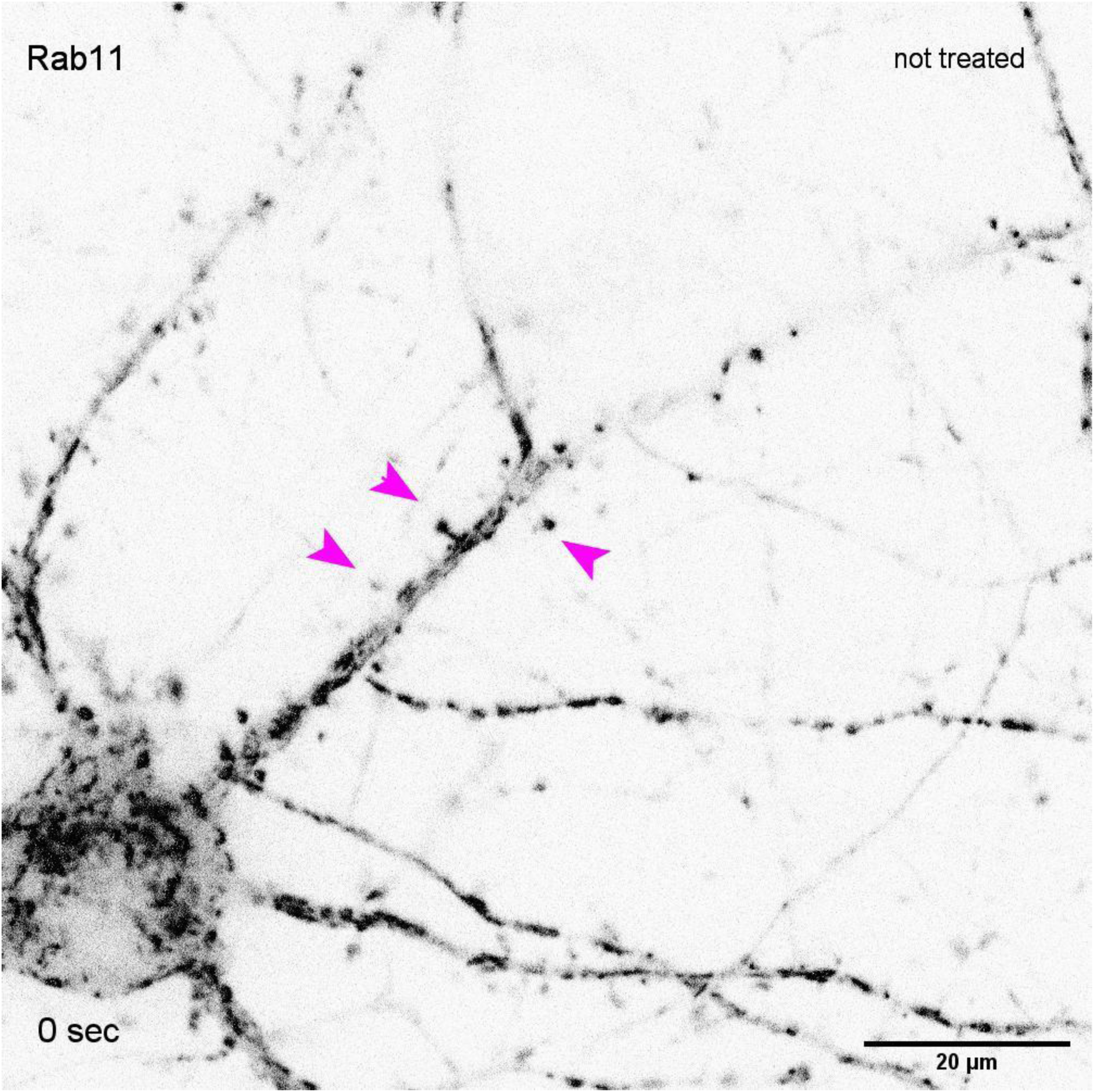
GFP-Rab11a channel of the same neuron as in Movie 1. Arrows point to examples of spines where mobile vesicles are present.

**Movie 3.**
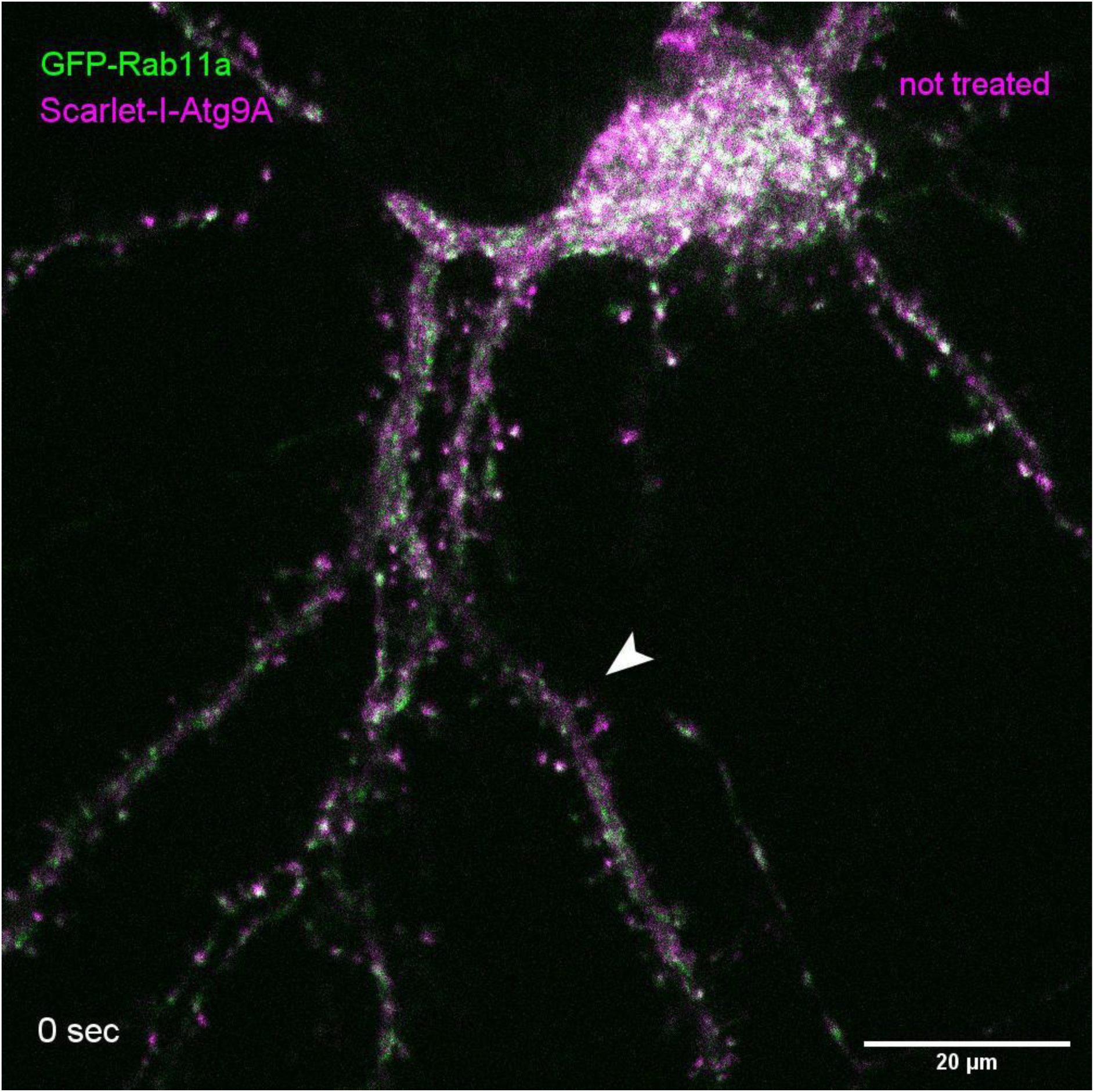
Hippocampal neuron transfected with plasmids encoding GFP-Rab11a (green) and Scarlet-I-Atg9A (magenta) over 1 min. Arrows point to mobile Atg9A.

**Movie 4.**
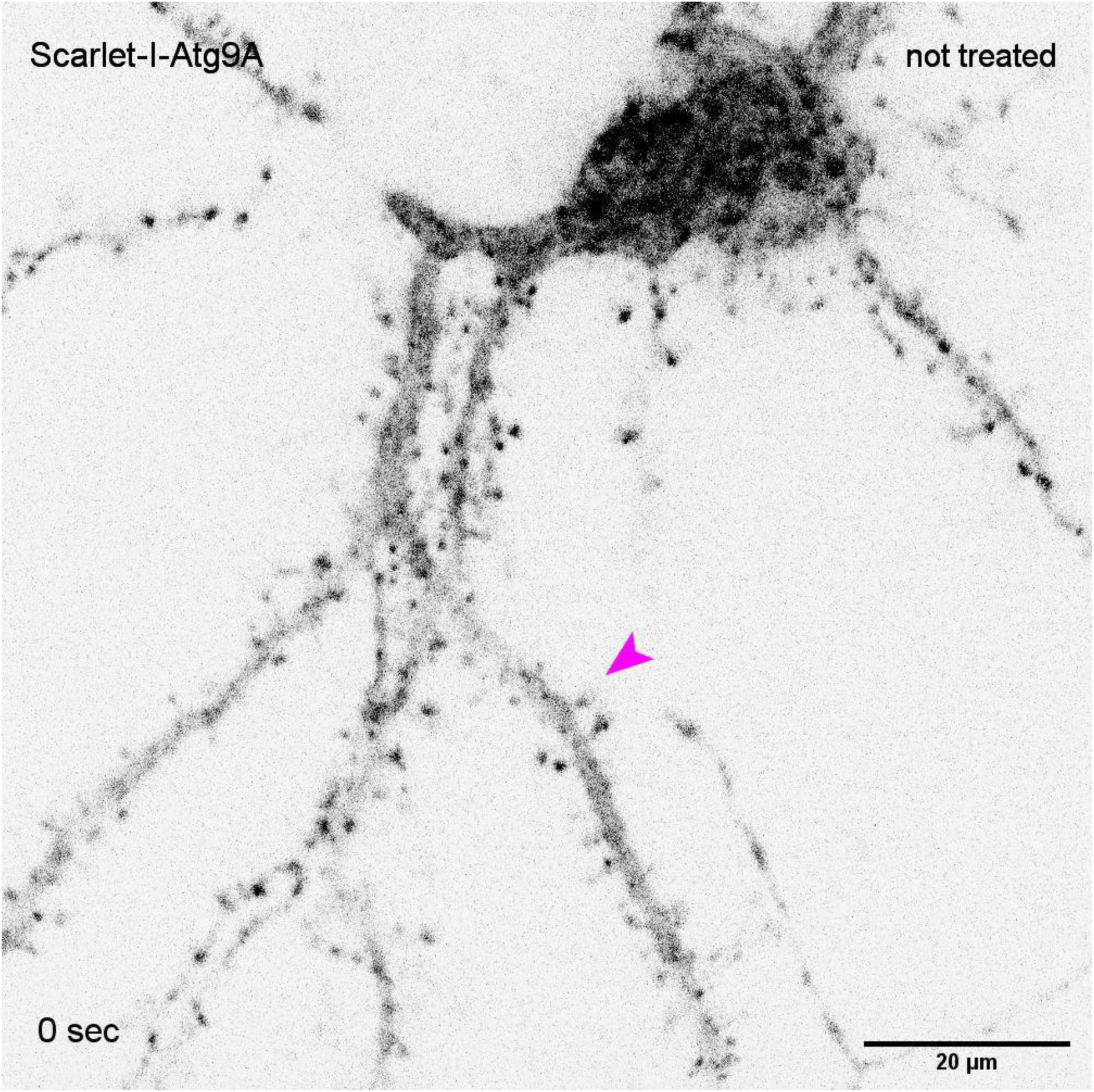
Atg9A channel of the same movie as in Movie 3. Arrow points to mobile Atg9A.

**Movie 5.**
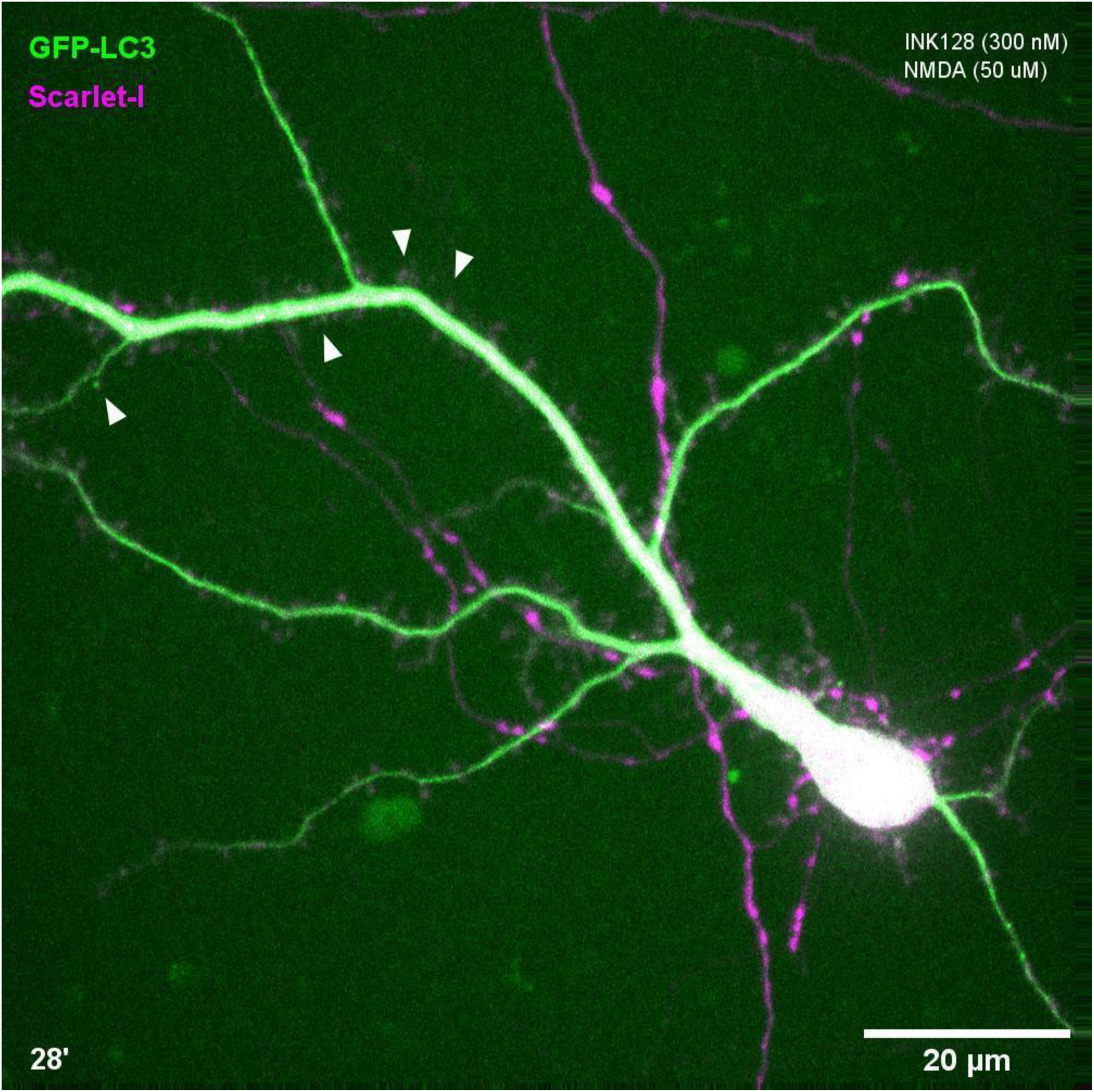
Hippocampal neuron transfected with EGFP-LC3 (green) and Scarlet-I (magenta) upon Ink128+NMDA treatment over 44 min. Arrows point to the emerging EGFP-LC3-positive puncta.

**Movie 6.**
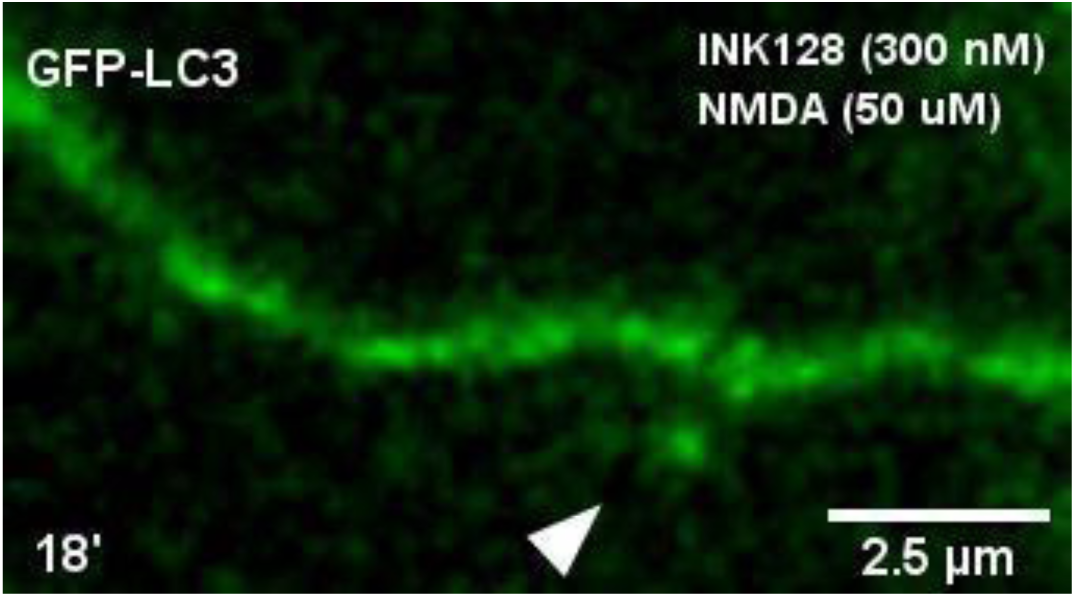
Fragment of a dendrite of a hippocampal neuron transfected with plasmid encoding EGFP-LC3 (green) and Scarlet-I (magenta) upon INK128+NMDA treatment over 44 min. Arrow points to the emerging EGFP-LC3-positive vesicle. EGFP-LC3 channel is shown.

**Movie 7.**
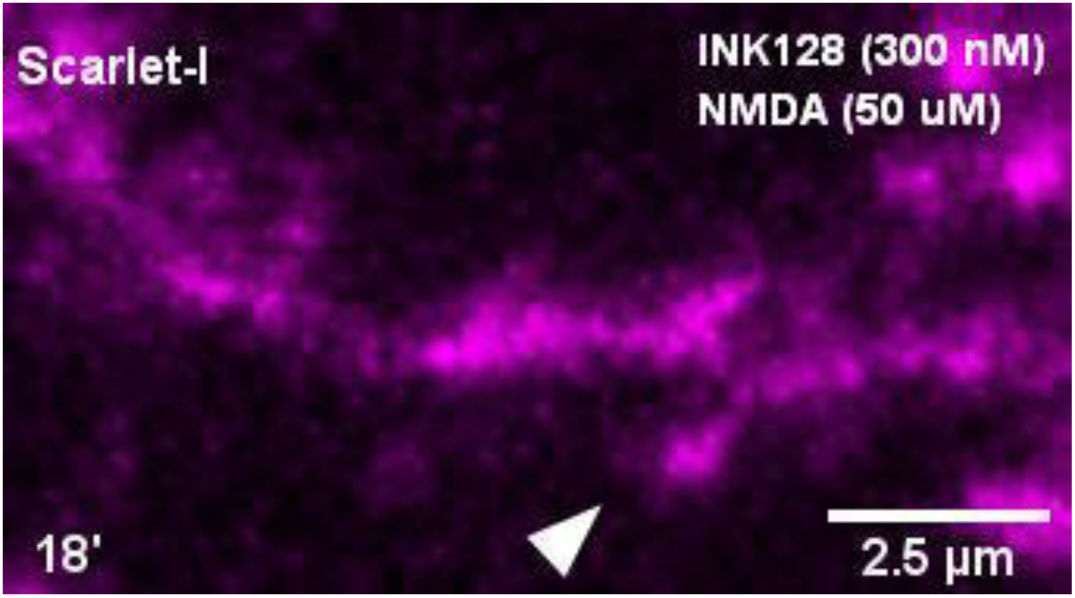
Fragment of the same dendrite as in Movie 6, Scarlet-I channel.

**Movie 8.**
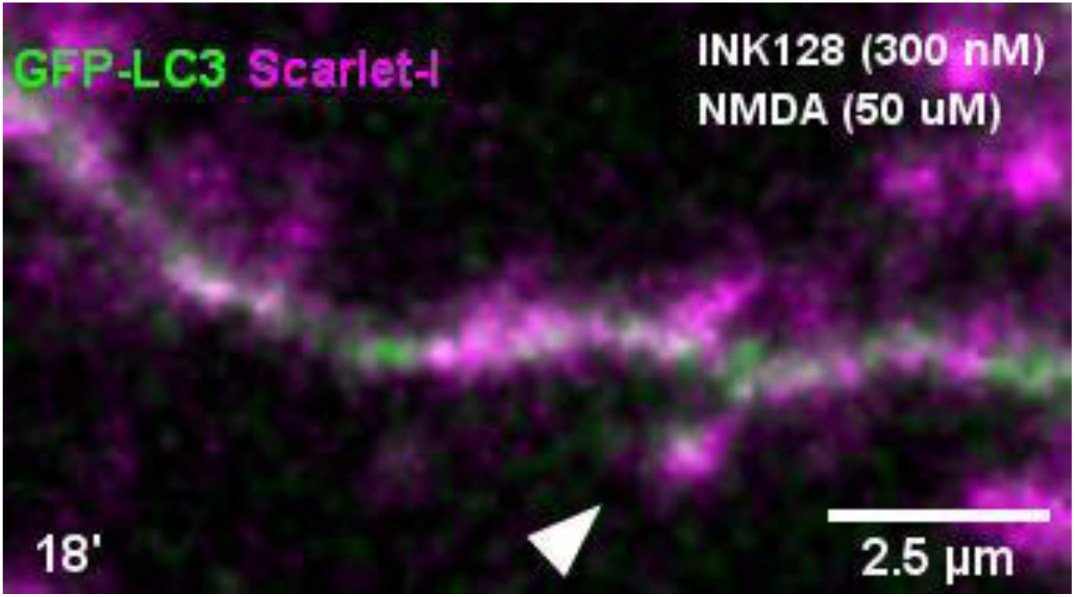
Fragment of the same dendrite as in Movie 6, both channels.

